# Co-Targeting IL-6 and EGFR signaling for the treatment of schwannomatosis and associated pain

**DOI:** 10.1101/2023.02.06.527377

**Authors:** Zhenzhen Yin, Limeng Wu, Yanling Zhang, Yao Sun, John W. Chen, Sonu Subudhi, William Ho, Grace Y. Lee, Athena Wang, Xing Gao, Jun Ren, Chao Zhu, Na Zhang, Gino B. Ferraro, Alona Muzikansky, Luo Zhang, Anat Stemmer-Rachamimov, Jianren Mao, Scott R. Plotkin, Lei Xu

**Affiliations:** Edwin L. Steele Laboratories, Department of Radiation Oncology, Massachusetts General Hospital and Harvard Medical School, Boston, Massachusetts 02114; Institute for Innovation in Imaging, Center for Systems Biology, and Division of Neuroradiology, Department of Radiology, Massachusetts General Hospital, Boston, Massachusetts 02114; St. Mark’s School, Southborough, MA, 01772; Department of Biomedical Engineering, Boston University College of Engineering, Boston, MA, 02215; Division of Biostatistics, Massachusetts General Hospital, Harvard Medical School, Boston, MA, 02114; Department of Otolaryngology - Head and Neck Surgery, Beijing TongRen Hospital, Capital Medical University, Beijing 100730, China; Molecular Pathology Division, Massachusetts General Hospital, Boston, MA, 02114; Center for Translational Pain Research, Massachusetts General Hospital, MA 02114; Department of Neurology and Cancer Center, Massachusetts General Hospital, Boston, MA, 02114

## Abstract

Patients with Schwannomatosis (SWN) overwhelmingly present with intractable, debilitating chronic pain. There are no effective therapies to treat SWN. The drivers of pain response and tumor progression in SWN are not clear. The pain is not proportionally linked to tumor size and is not always relieved by tumor resection, suggesting that mechanisms other than mechanical nerve compression exist to cause pain. SWN research is limited by the lack of clinically-relevant models. Here, we established novel patient-derived xenograft (PDX) models, dorsal root ganglia (DRG) imaging model, and combined with single-cell resolution intravital imaging and RNASeq, we discovered: i) schwannomas on the peripheral nerve cause macrophage influx into the DRG, via secreting HMGB1 to directly stimulate DRG neurons to express CCL2, the key macrophage chemokine, ii) once recruited, macrophages cause pain response via overproduction of IL-6, iii) IL-6 blockade in a therapeutic setting significantly reduces pain but has modest efficacy on tumor growth, iv) EGF signaling is a potential driver of schwannoma growth and escape mechanism from anti-IL6 treatment, and v) combined IL-6 and EGFR blockade simultaneously controlled pain and tumor growth in SWN models. Our findings prompted the initiation of phase II clinical trial (NCT05684692) for pain relief in patients with SWN.

## Main

Schwannomatosis (SWN) is a genetic disorder characterized by multiple non-malignant schwannomas growing on the spine and peripheral nerves ^1–3^. Patients with SWN overwhelmingly present with intractable, chronic pain, which can be severe enough to cause permanent disability ^4–7^. The etiology of pain in SWN is not clear. There is no single pattern of pain reported by patients. In some cases, pain is associated with a specific schwannoma and resolves after surgical excision. However, in many patients, pain can recur after surgical excision, even without recurrence of the schwannoma. In other patients, pain is multifocal or generalized and may not correspond to any pattern of schwannomas. These suggest that additional mechanisms other than mechanical nerve compression exist to cause pain. There are no effective therapies that halt tumor growth or ameliorate SWN-associated pain ^8^. Treatment for SWN is limited to surgery, which carries a major risk of iatrogenic nerve injury ^9^. SWN patients with chronic pain are typically treated with a variety of conventional pain medications, including anti-inflammatory drugs, neuropathic pain medications, and opioids. However, a majority of patients continue to rate their pain as moderate to severe despite these aggressive options ^8–10^. The suffering and disability, in combination with the lack of FDA-approved pharmacologic treatment options, make the effective treatment of SWN a major unmet medical need.

As the drivers of pain response and tumor progression in SWN remain obscure, the development of effective treatments for SWN and related pain has been extremely slow and inefficient. One of the biggest obstacles in SWN research is the lack of clinically-relevant models - no SWN cell lines are available, and as a non-malignant tumor, no patient-derived xenograft (PDX) models are available. Here, we report the establishment of i) schwannoma cell lines from SWN patients with varying degrees of pain, ii) orthotopic sciatic nerve and spine PDX modeling schwannomas growing on the peripheral nerves and on the spine in patients and faithfully reproducing tumor-induced pain, and iii) dorsal root ganglia (DRG) imaging model that allows real-time, longitudinal imaging of the DRG microenvironment, which contains the primary sensory neurons and relays pain signals from the peripheral nerve into the central nervous system. Using these novel tools, we deciphered the cellular and molecular crosstalk between schwannoma (HMGB1)– neuron (CCL2)–macrophage (IL-6) in driving pain response, and identified EGF pathway as driver of SWN tumor progression.

## Results

### Establish orthotopic PDX SWN models

We established 9 patient-derived SWN cell lines using surgically resected schwannomas tissues from SWN patients with varying degrees of pain - four non-painful tumors and five painful tumors (Supplementary Table S1). SWN grows on the peripheral nerves (in 89% of patients) and spine (in 74% of patients) ^1^’^6^. Therefore, we established orthotopic sciatic nerve and spine models in nude mice. In the sciatic nerve model, we used 3D ultrasound to confirm tumor formation and location relative to the sciatic nerve (Fig 1A). We observed that sham surgery-induced acute pain returned to normal levels 5-7 days post-surgery; with tumor growth, pain response persisted (Fig 1B-C), indicating tumor-induced pain response.

**Figure 1.**
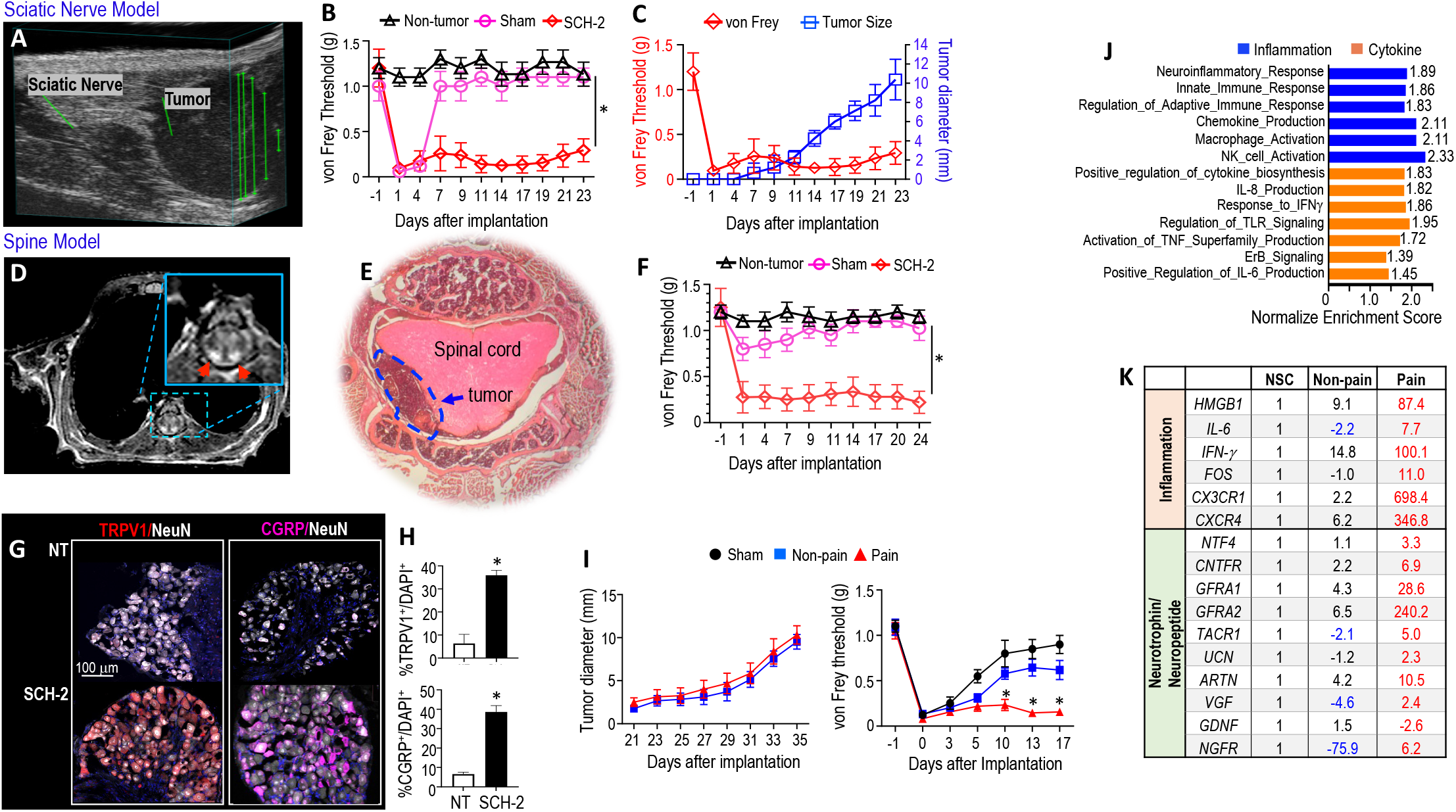
Establishment and characterization of orthotopic PDX SWN models. In the sciatic nerve model **(A)** 3D ultrasound showed a predominately hypoechoic ovoid mass consistent with a schwannoma abutting and exerting mild mass effect on the sciatic nerve. **(B)** von Frey filament test was performed at baseline and at the indicated time points post-implantation. **(C)** Changes of von Frey threshold and tumor diameter over time. In the spine model **(D)** Post-contrast MR images demonstrated confluent contrast-enhancing intradural extramedullary masses abutting the posterior spinal cord (red arrows), shown on the axial plane, and **(E)** H&E staining revealed mild compression of the spinal cord by the tumor (arrow, circled area). (**F)** von Frey filament test was performed at baseline and at the indicated time points post-implantation. Non-tumor and sham groups, (n=4/ea), SCH2 (n=12). *P<0.0001. **(G)** Representative IF staining images of nociceptive TRPV1 (red) and CGRP (pink) in DRG neurons (NeuN, white) in non-tumor bearing mice (NT) and in SCH2 tumor-bearing mice. **(H)** The percent of positively stained cells was manually counted. N=12 DRGs/ea, *P<0.0001. **(I)** von Frey filament test and tumor growth curve comparison between non-painful SCH-1 with painful SCH-2 tumors in the sciatic nerve model. N=8/each. *P<0.001. **(J)** Normalized enrichment scores from bulk RNASeq showing pathways elevated in patient-derived schwannomas (SCH-1, SCH-2, and SCH-4) compared to normal Schwann cells (n=3). **(K)** Differentially expressed genes in painful SCH-2 tumor vs. non-pain SCH-1 tumor, compared to normal Schwann cells (NSC). All animal studies are presented as mean±SEM, and are representative of at least three independent experiments.

In the spine model, we used MR imaging and H&E staining to confirm that schwannoma xenografts grew in the intradural and extramedullary space of the spine (Fig 1D-E), reproducing the correct anatomical location of spinal schwannomas in patients. As intrathecal injection does not require surgery, sham injection only induced a mild pain response in the spine model that returned to normal level within 3 days; in the PDX model, mechanical allodynia increased (Fig 1F).

In both sciatic nerve and spine models, no differences in tumor-induced pain response were observed between male and female mice (Fig S1A), and tumor growth did not affect motor function in mice during the course of pain response measurement as evaluated by rotarod test (Fig S1B). Histological analysis confirmed the Schwann cell origin and schwannoma histology (Fig S1C-D). Immunofluorescent (IF) staining further confirmed that, in both models, tumor growth induced the expression of nociception-related molecules in DRG sensory neurons, including transient receptor potential vanilloid 1 (TRPV1) and calcitonin gene-related peptide (CGRP)^11^(Fig 1G-H).

Next, we examined if our PDX model could reproduce the pain status of the tumor from SWN patients. We implanted the SCH-1 cells from a non-painful left accessory nerve schwannoma and the SCH-2 cells from a painful right pudendal nerve schwannoma in the sciatic nerve model. The non-painful tumors and painful tumors grew at the same rate; however, the painful SCH-2 tumors elicited a significantly greater pain response compared to the non-painful SCH-1 tumors (Fig 1I). These studies demonstrate that our model can faithfully reproduce tumor-induced pain response in SWN.

### Inflammatory gene signature in painful schwannomas

To elucidate the molecular mechanisms of tumor-induced pain response, we used RNA sequencing (RNASeq) to compare the transcriptome of patient schwannomas with normal Schwann cells. Pathway enrichment analysis was performed on 487 genes that were differentially expressed (FDR q-value<0.05 and Log2FC>2, Supplementary Table S2). Gene Set Enrichment Analysis showed that in schwannomas, elevated genes are significantly enriched in the following pathways: i) neuroinflammatory response, ii) inflammatory cell activation, such as macrophages and NK cells, and iii) production of inflammatory cytokines, such as TNFa, IL-6, and IFN-g (Fig 1J).

We further identified a set of genes significantly elevated in painful tumors compared to non-painful tumors. This panel of pain signature genes includes inflammatory cytokines, neurotrophin, and neuropeptide (Fig 1K). These profiling data prompted us to explore possible mechanisms governing neuroinflammation-induced pain in SWN.

### Peripheral nerve schwannoma triggers macrophage influx into the DRG

In mice bearing SCH2 tumor in the sciatic nerve, the DRG sensory neurons express significantly higher levels of C-C motif ligand 2 (CCL2), the key macrophage chemokine (Fig 2A), suggesting recruitment of macrophage. Macrophages play a critical role in inflammation ^12^. To track macrophage infiltration into the DRG, we custom-designed and 3D-printed a DRG imaging window and performed intravital imaging to directly observe tumor-induced changes in the DRG microenvironment, in real-time at single-cell resolution throughout the course of tumor formation. C-C chemokine receptor 2 (CCR2), the main receptor for CCL2, is expressed primarily on macrophages ^12^. To visualize macrophages, we used *Ccr2^RFP/RFP^* reporter mice. However, the reporter mice are in the immune-competent C57/BL6 background. To remedy this and grow the patient-derived tumors, we injected bone marrow-derived macrophages (BMM) from *Ccr2^RFP/RFP^* mice into whole-body irradiated nude mice. After the successful generation of bone marrow chimera, we implanted the imaging window over the DRG and waited 7 days to minimize window implantation-induced inflammation and pain before tumor implantation and imaging (Fig 2B). Intravital imaging of the DRG revealed a 5.3 ± 0.92 fold increase in *Ccr2^RFP^* macrophages one day after tumor implantation; this increase was sustained until day 14 post-tumor implantation, while non-tumor bearing mice had few to no RFP-expressing macrophages in the lumbar DRG (Fig 2C-D). The induction of macrophage infiltration into the DRG was confirmed by flow cytometry (Fig 2E) and histological analysis (Fig S2) - in the lumbar DRGs obtained from tumor-bearing mice, there are significantly more macrophages and skewed toward the inflammatory M1 phenotype as compared to DRGs from non-tumor bearing mice.

**Figure 2.**
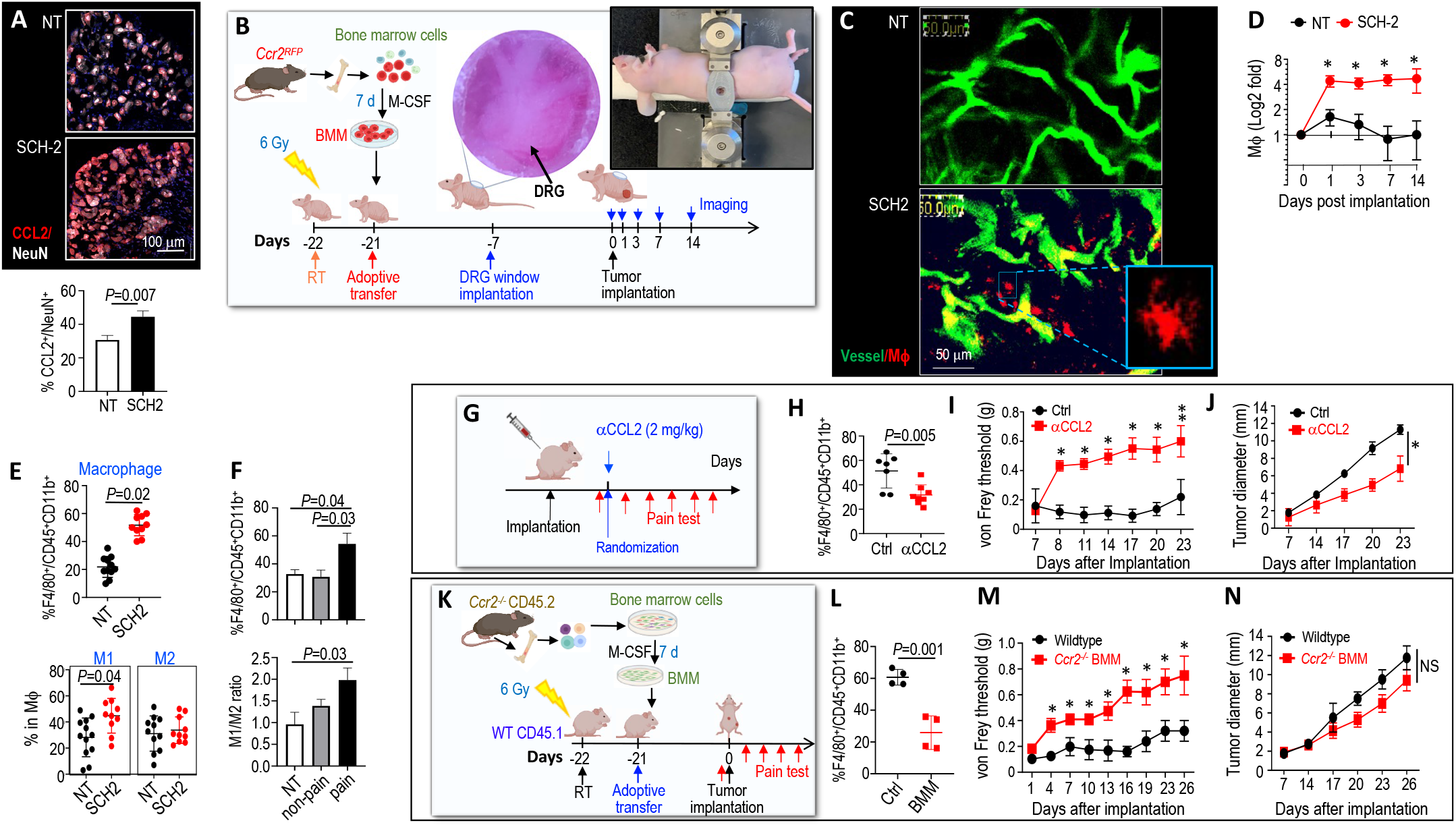
Peripheral nerve schwannomas trigger macrophage influx into the DRG to mediate pain response. **(A)** Representative IF staining images of CCL2 (red) in the lumbar DRG neurons (NeuN, white) collected from non-tumor bearing mice and mice bearing SCH-2 tumors in the sciatic nerve (ipsilateral L3-L6 DRGs). The percent of positively stained cells was manually counted. N=12 DRGs/ea. **(B)** Schematic of bone marrow-derived macrophage adoptive transfer, DRG window implantation, sciatic nerve SCH2 tumor implantation, and imaging timeline. **(C)** Representative 2-photon images of DRG vessels (green) and infiltrating *Ccr2^RFP^* macrophages (red). **(D)** Fold change of macrophages in both groups at different times post-tumor implantation. *P<0.005, n=4 mice/group. Flow cytometry analysis of the number and the phenotype of macrophages in lumbar DRGs (L3-L6) ipsilateral to sciatic nerve tumor between **(E)** non-tumor bearing (NT, n=12) vs. painful SCH-2 tumors (n=10), and **(F)** non-painful SCH-1 vs. painful SCH-2 tumors. N=5 mice/ea. **(G)** Schematic of αCCL2 treatment in the SCH-2 sciatic nerve model. **(H)** Flow cytometry analysis of macrophages in the lumbar DRGs. **(I)** von Frey filament test. *P<0.001, **P=0.03. **(J)** Tumor diameter as measured by caliper. *P=0.01. Control, n=7; αCCL2, n=8. **(K)** Schematic depicting the timeline of *Ccr2^-/-^* macrophage adoptive transfer, SCH2 tumor sciatic nerve implantation, and pain test. **(L)**Flow cytometry analysis of macrophages in the lumbar DRGs. **(M)**von Frey filament test. *P<0.001. **(J)** Tumor diameter as measured by caliper. N=11 mouse/group. All animal studies presented are mean±SEM, and represent at least three independent experiments. Flow cytometry data are presented as mean±SD.

### Macrophage in the DRG is essential in tumor-induced pain

To characterize the functional role of DRG macrophages in schwannoma-induced pain response, we first compared the macrophage infiltration in painful vs non-painful models. Compared to the non-painful SCH-1 model and non-tumor bearing mice, mice bearing painful SCH-2 tumors have significantly more macrophages in the DRG, and these macrophages are in the M1 pro-inflammatory phenotype (Fig 2F), suggesting a role of the macrophage in the pain phenotype.

To determine if the DRG macrophages are essential in SWN-induced pain response, we blocked macrophage infiltration by anti-CCL2 antibody treatment (αCCL2, Fig 2G). αCCL2 treatment: i) successfully reduced macrophage infiltration into the DRG (Fig 2H), ii) significantly reduced pain in the mice (Fig 2I), and iii) inhibited tumor growth (Fig 2J). CCL2 can directly stimulate neurons in the pain cascade ^13^. Therefore, to dissect the contribution of macrophage from the CCL2 direct effect on neurons, we performed adoptive transfer of bone marrow-derived macrophages from *Ccr2*^-/-^ mice (Fig 2K). In the chimera mice, *Ccr2*^-/-^ macrophages exhibit defective chemotaxis into the DRG (Fig 2L), schwannomas grow at the same rate but induce less pain compared to the wild-type mice (Fig 2M-N). These studies reveal that macrophages are essential in mediating tumor-induced pain in SWN models.

### Schwannoma-secreted HMGB1 upregulates CCL2 in DRG neurons

To identify the driver of macrophage recruitment, we examine the transcriptome profiling data and discovered that painful tumors express higher levels of HMGB1 (Fig 1K). HMGB1 is a potent inflammation initiator and amplifier and has been shown to upregulate CCL2 in smooth muscle cells in an artery injury model ^14^. We confirmed that patient-derived schwannoma cells express elevated levels of HMGB1 in vitro (Fig S3), and both sciatic nerve and spine tumors secrete HMGB1 into the circulation (Fig 3A). Using an *ex vivo* DRG slice culture model, SWN tumor cell-conditioned medium (TC-CM) significantly stimulated CCL2 expression in DRG neurons; and application of HMGB1 inhibitor - Glycyrrhizin - abolished the tumor-induced neuronal CCL2 expression (Fig 3B-C), suggesting that HMGB1 regulates neuronal CCL2 expression.

**Figure 3.**
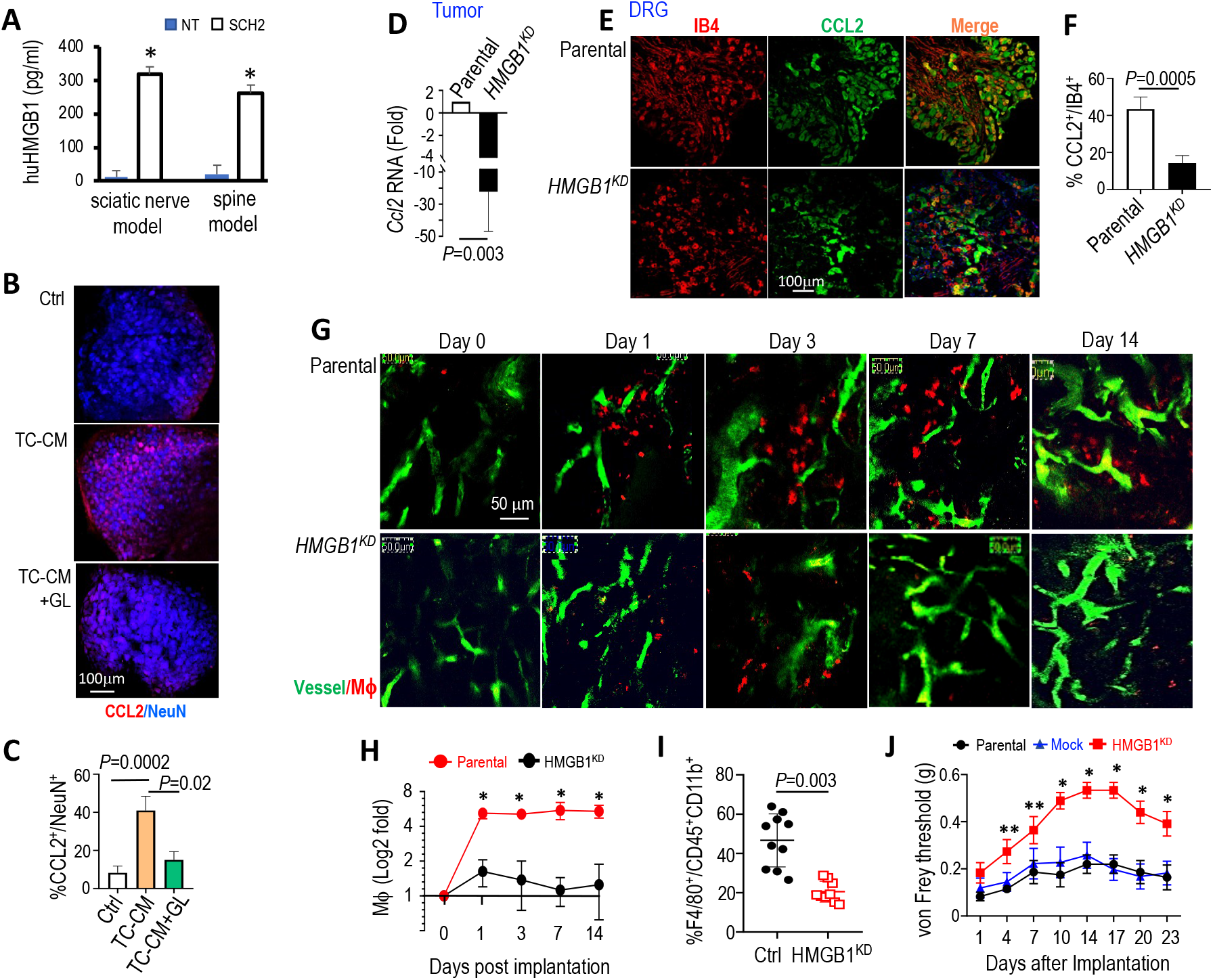
Schwannoma-secreted HMGB1 regulates CCL2 expression in DRG neurons and macrophage recruitment. **(A)** ELISA measurement of the human HMGB1 protein in the plasma of non-tumor bearing mice (NT) and SCH-2 sciatic nerve and spine models. N=4 mice/ea. *P<0.001. **(B)** Lumbar DRGs were collected from nude mice and cultured in DMEM (Ctrl), SCH-2 cell conditioned medium (TC-CM), or TC-CM+HMGB1 inhibitor, Glycyrrhizin (GL, 50 μg/ml). 48 hours later, DRG slices were fixed and IF stained for CCL2 (red) in DRG neurons (NeuN, blue). **(C)** Quantification of the % neurons that express CCL2 using ImageJ. N=3 DRG slices/group. Mice were implanted with parental, mock and *HMGB1^KD^* cells in the sciatic nerve: **(D)** At the experiment endpoint (tumor reaches 1 cm in diameter), tumor tissues were collected for qRT-PCR analysis of mouse *Ccl2* mRNA level change. N=3/group and **(E)** Lumbar DRGs (L3-6) ipsilateral to the sciatic nerve tumors were collected and stained for CCL2 (green) in DRG neurons (IB4, red). **(F)** The percent of positively stained cells was manually counted. N=12 DRGs/ea. **(G)** Representative 2-photon images of macrophage (red) infiltrating the DRG at different timepoint after tumor implantation. DRG vessels (green). **(H)** Fold change of macrophages in both groups at different times post-tumor implantation. *P<0.001, n=4 mice/group. **(I)** Flow cytometry analysis of macrophages in the lumbar DRGs. N=12 DRGs/group. **(J)** von Frey filament test. N=8 mouse/group. *P<0.05, **P<0.001. All animal studies are presented as mean±SEM, and are representative of at least three independent experiments. All expression analysis and flow data are presented as mean±SD.

To investigate if HMGB1 plays a direct causal role in neuronal CCL2 expression, we genetically silenced HMGB1 using CRISPR-Cas9 gene editing. Single-cell clones were picked after antibiotic selection and three clones with the lowest HMGB1 level were pooled to avoid clonal variation (Fig S3B). HMGB1 knockdown did not affect tumor cell viability *in vitro* nor tumor growth in the PDX model (Fig S3C-D). HMGB1 knockdown significantly reduced CCL2 expression in the tumors (Fig 3D) and in the DRG sensory neurons (Fig 3E-F). Using DRG intravital imaging, we observed that in mice bearing parental tumors, macrophages infiltrate the DRG as early as Day 1 post-implantation and persisted until Day 14; whereas in mice bearing the HMGB1-knockdown tumors, significantly fewer macrophages were observed in the DRG (Fig 3G-H). The imaging finding was confirmed by flow cytometry analysis of the lumbar DRG macrophages (Fig 3I). Consistent with the reduced macrophage infiltration, mice bearing HMGB1 knockdown tumors exhibited significantly reduced pain (Fig 3J).

### αIL6 treatment reduces pain

To decipher how tumor affects macrophages to trigger pain response, we differentiated macrophages (Raw264.7 cell) to different phenotypes by treating them with: i) tumor cell-conditioned medium (TC-CM) - representing TAMs, ii) regular culture medium - representing naïve M0 phenotype, and iii) IFN-γ (1000 U/ml) - inducing an M1 inflammatory phenotype and serving as a positive control for nociception-related signaling ^15,16^. We collected CM from these macrophage cultures and added them to the organotypic DRG slice culture. After 3 days, we found that CM from TAMs, as well as from the M1 inflammatory macrophages, significantly induced nociceptive TRPV1 and CGRP expression in DRG neurons (Fig 4A). Analyzing the macrophage-secreted factors, we found that TC-CM significantly induces the secretion of proinflammatory cytokines IL-6, IL-1ß, and TNFα in macrophage Raw264.7 cells and in mouse peritoneal macrophages (Fig 4B).

**Figure 4.**
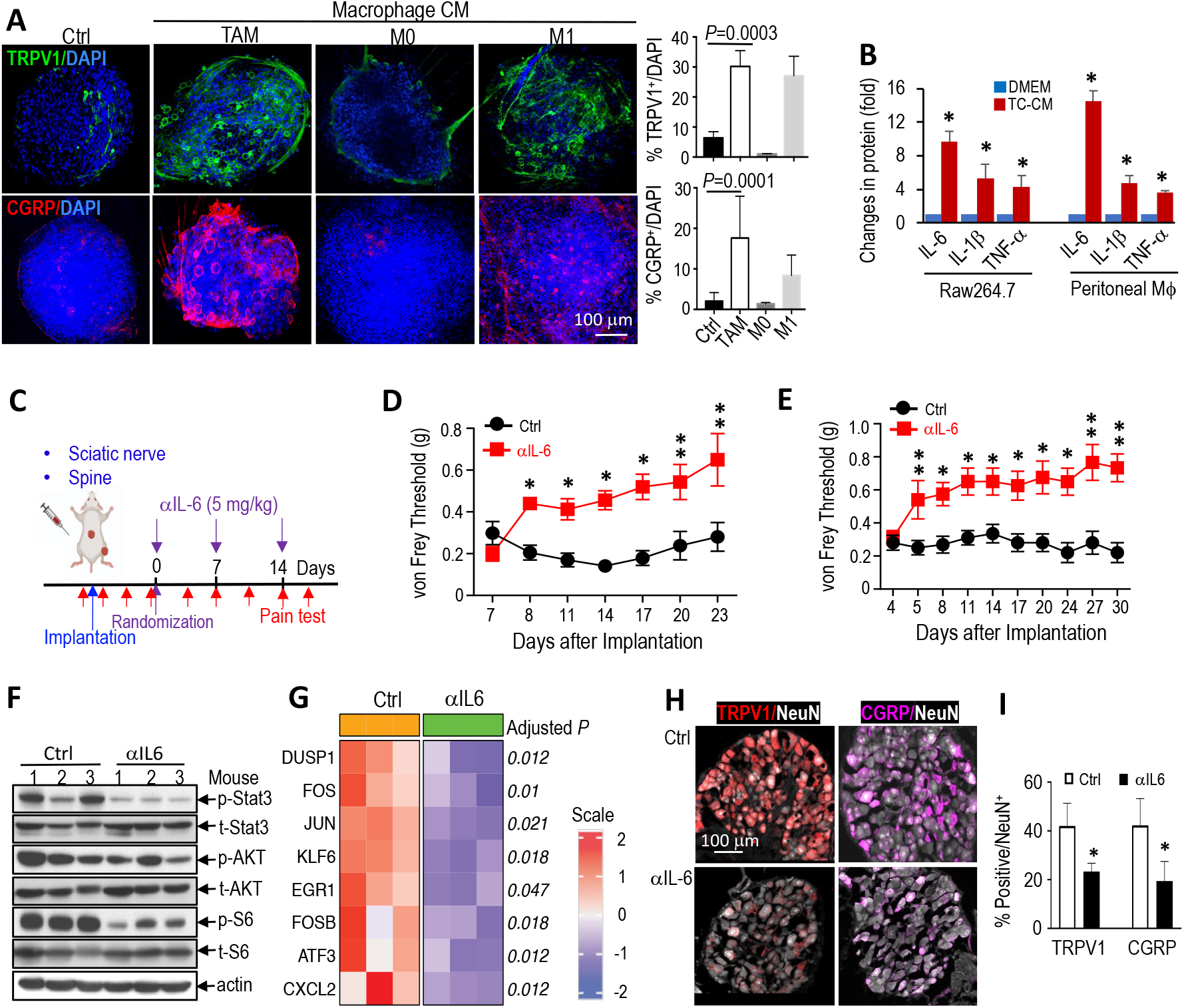
αIL6 reduces pain. **(A)** Representative images of IF staining for TRPV1 (green) and CGRP (red) in the DRG slice culture model. Quantification of the percent positive staining was analyzed by ImageJ. DAPI, blue. N=3 DRG slices/group. **(B)** ELISA of murine IL-6, IL-1ß, and TNF-α protein in the supernatant of Raw264.7 cells and peritoneal macrophages cultured in DMEM or TC-CM for 48 hours. *P<0.001. **(C)** Schematic and timeline of αIL-6 treatment experiment. von Frey filament threshold in **(D)** the sciatic nerve model. N=10/group. *P<0.0001, **P<0.05, and **(E)** the spine model. N=11/group. *P<0.002, **P<0.05. **(F)** Western blot of tumor tissues, n=3 mice/group. **(G)** Heatmap showing top differentially expressed genes between TAMs sorted from αIL6 and control mice. Adjusted P value (Benjamini & Hochberg method) depicted on the right side of the heatmap. **(H)** Representative IF staining images for nociceptive TRPV1 (red) and CGRP (pink) in neurons (NeuN, white) of L3-6 DRGs ipsilateral to sciatic nerve tumor in control and αIL-6 treated mice. **(I)** The percent of positively stained cells was manually counted. N=12 DRGs/ea, *P<0.0001. All animal studies are presented as mean±SEM and are representative of at least three independent experiments. All expression analyses are presented as mean±SD.

IL-6, the most significantly induced pro-inflammatory cytokine in tumor-primed macrophages, is a major trigger for inflammatory pain in a variety of pain models ^17,18^. In both sciatic nerve and spine models, blocking IL-6 signaling in a therapeutic setting with anti-IL-6 (αIL-6) significantly alleviated pain response (Fig 4C-E). Mechanistically, αIL-6 treatment reduced inflammatory STAT3 and AKT/S6 signaling in tumor tissues (Fig 4F). We sorted TAMs and performed RNASeq, we observed that genes related to inflammation are significantly reduced in αIL-6-treated tumors (Fig 4G). In accordance with these reduced inflammatory responses, we observed that αIL-6 treatment reduced the expression of nociceptive TRPV1 and CGRP in the DRG sensory neurons (Fig 4H-I)

### Combined IL-6 and EGFR blockade concurrently control tumor and pain

Although αIL-6 treatment reduced pain response, it only modestly delayed tumor growth (Fig 5A). We investigated potential mechanisms for the low tumor control efficacy and found that EGFR and ErbB2/3 phosphorylation increased in the αIL-6 treated tumors, suggesting pathway activation (Fig 5B). In patient-derived schwannoma cells, we also detected elevated baseline EGFR, ErbB2, and ErbB3 phosphorylation as compared to normal Schwann cells (Fig 5C). These data suggest that EGF signaling may be a driver of schwannoma progression and escape mechanism from αIL-6 treatment.

**Figure 5.**
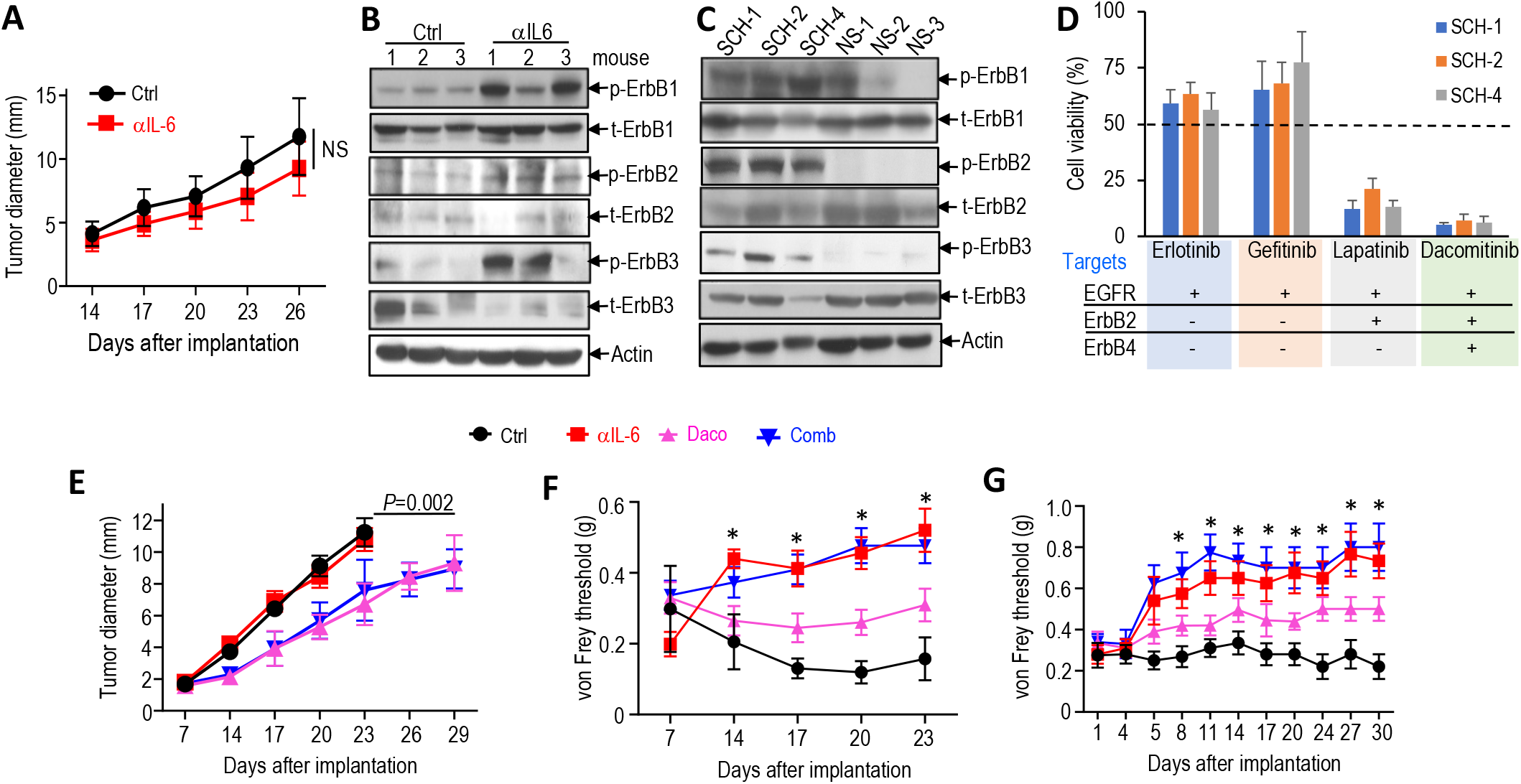
Combined IL-6 and EGFR blockade concurrently control tumor and pain. **(A)** Tumor size was measured by caliper every 3 days in the sciatic nerve model. Ctrl N=8, αIL-6 N=10. Western blot of total- and phosphorylated-ErbB1, ErbB2, and ErbB3 in **(B)** control and αIL-6 treated tumors and in **(C)** SCH1, −2, −4 cells and normal Schwann cells (NS) lysates. **(D)** In vitro drug screening by MTT assay. Cells were treated with DMEM or 10 μM of each inhibitor for 72 hours. Data are presented as mean±SD. **(E)** Tumor growth curve, and **(F)** von Frey filament threshold in the sciatic nerve model, N=12/group. **(G)** von Frey filament threshold in the spine model. N=12/group. *P<0.0001 for both comparisons between αIL-6 vs. Ctrl and combination (comb) vs. Ctrl. All animal studies are presented as mean±SEM, and are representative of at least three independent experiments.

To test the treatment effects of EGFR blockade, we first performed *in vitro* screening of four FDA-approved EGFR tyrosine kinase inhibitors (TKIs) ^19^. Among the 4 inhibitors, lapatinib (an EGFR and HER2 TKI) and dacomitinib (a pan-ErbB inhibitor) significantly reduced the viability of patient-derived schwannoma cells, while erlotinib and gefitinib (EGFR inhibitors) had minimal effects (Fig 5D). We focused on testing the treatment potential of dacomitinib, which elicited the most effective inhibition on cell growth *in vitro*.

In the PDX model, dacomitinib treatment alone or in combination with αIL-6 significantly delayed tumor growth as compared to control or αIL-6 alone, however, combination treatment did not offer additive effects compared to dacomitinib monotherapy (Fig 5E). In both sciatic nerve and spine models, dacomitinib monotherapy had minimal effects on reducing pain; αIL-6 alone or in combination with dacomitinib significantly reduced pain; however, no additive effects were observed from the combination treatment as compared to αIL-6 alone (Figs 5F-G). None of the treatments caused systemic toxicity as evaluated by body weight loss (Fig S4A). In the tumor tissue, we found that dacomitinib alone and in combination with αIL-6 significantly reduced tumor cell proliferation and increased tumor cell apoptosis (Fig S4B).

### Elevated inflammation in patients with SWN

To prepare the translation of our study into the clinic, we validated our pre-clinical findings in archived human SWN tumor samples. Compared to the normal nerve, we found a trend of elevated infiltrating TAMs, a significantly elevated CCL2, HMGB1, IL-6 expression, and STAT3 and EGFR phosphorylation (Fig 6). These studies confirm that IL-6 and the EGF pathway are valid targets in SWN.

**Figure 6.**
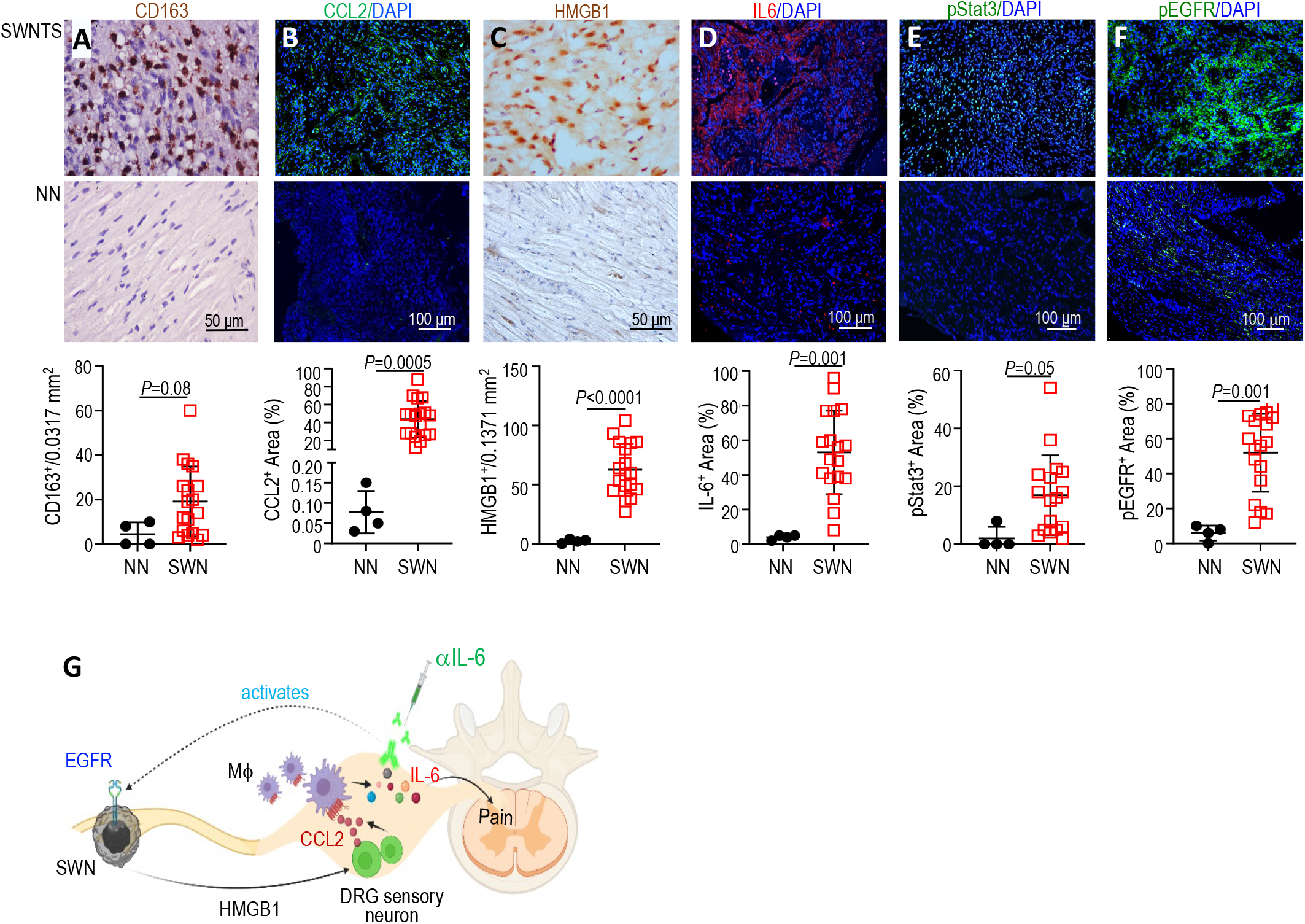
Elevated inflammation in patients with SWN. Archived paraffin-embedded tumors from patients with SWN were immunostained for **(A)** macrophage (CD163^+^, brown), **(B)** CCL2 (green), **(C)** HMGB1 (brown), **(D)** IL-6 (red), **(E)** phosphorylated STAT3 (green), and **(F)** phosphorylated EGFR (green). The number of TAMs and HMGB1 positively stained cells per 0.0317mm^2^ field was counted manually. Fluorescent images were analyzed for the % positive stained area using ImageJ software. **(G)** Schwannoma-secreted HMGB1 stimulates DRG sensory neurons to express CCL2, which recruits macrophages into the DRG. Macrophages cause pain response via overproduction of IL-6, and combined IL-6 and EGFR blockade can concurrently reduce pain and inhibit tumor growth in SWN models.

## Discussion

In this study, using novel PDX models and single-cell resolution intravital imaging, we identified drivers for both tumor growth and pain in SWN models. We discovered that schwannomas grew in the peripheral nerve, via secreting HMGB1 into the circulation, stimulate DRG sensory neurons to express CCL2 to recruit macrophages. Macrophages are essential in mediating pain via overproducing IL-6. Blocking IL-6 signaling in a therapeutic setting reduced tumor-induced pain but had modest efficacy in tumor control in the PDX models. We identified EGF signaling as a driver of schwannoma growth and a potential escape mechanism from anti-IL6 treatment. We demonstrated that combined IL-6 and EGFR blockade simultaneously controlled pain and tumor growth in SWN models (Fig 6G).

One of the biggest challenges in basic science and translational SWN research is the lack of clinically-relevant models. Germline mutations in *SMARCB1* and *LZTR1* are known to cause SWN ^2^. Genetically engineered mice (GEM) of SWN have been developed. Biallelic inactivation of *Smarcb1* and *Nf2* in the Schwann cell lineage develops tumorlets consisting of whorls of Schwann cells in the DRG that are reminiscent of schwannoma tumorlets in human patients ^20^. However, as the timing of tumor formation in GEM can be inconsistent and the percentage of mice that develop tumors can be small, it is difficult to perform reproducible, robust drug testing experiments with GEM. In the era of personalized medicine, patient-derived models are of utmost importance. However, as a non-malignant tumor syndrome, SWN has no reported PDX models. A previous study reported the isolation of patient-derived schwannoma cells and demonstrated that tumor cells secrete inflammatory cytokines and affect neuronal pain-associated gene expression *in vitro* ^21^. As a step further, we established two orthotopic PDX models mimicking human schwannomas in the peripheral nerve and spine. Both models faithfully reproduced schwannoma-induced pain. Further optimization of the PDX models, such as breeding the GEM into immunodeficient mouse strains with stromal cells, innate and adaptive immune cells all harboring the same gene mutations, could define additional avenues of host contribution to tumor-induced pain response. In addition, as SWN is a rare disease and only 2-5% of SWN patients surgically remove their schwannomas, future expansion of the SWN cell bank accumulating a sufficient number of cells from different patients will shed light on the correlation between gene mutations and the pain phenotype.

Real-time in vivo imaging allows the characterization of the tumor microenvironment. Here, we developed a DRG imaging window model and performed intravital imaging to directly observe tumor-induced changes in the DRG microenvironment, in real-time at single-cell resolution throughout the course of tumor formation. We discovered that schwannomas grew distantly on the peripheral nerve caused inflammatory macrophage influx into the DRG. This finding prompted us to ask two questions: i) how do tumor lesions in the peripheral nerve cause macrophage infiltration in the DRG, and ii) how do macrophages induce pain response in SWN models? We further combined the intravital imaging model with genetic knockdown studies and discovered a functional role of HMGB1 in regulating neuronal expression of chemokine that recruits macrophage into the DRG. Taken together, the orthotopic PDX models and the imaging model will help address a major bottleneck in the development of therapeutics for SWN, and provide the SWN research and clinical community with robust, and biologically diverse tools and information to explore new therapeutic targets to tackle this devastating disease.

We deciphered the cellular and molecular crosstalk between schwannoma (HMGB1) - Neuron (CCL2) - macrophage (IL-6) in generating pain. The findings presented here show that schwannomas secrete HMGB1 to directly regulate CCL2 expression in the DRG neurons, thus recruiting macrophages into the DRG to cause pain. These studies suggest that CCL2 and HMGB1 are also potential therapeutic targets for SWN. CCL2, a key macrophage chemokine, is implicated in tumor progression and metastasis ^22^, as well as directly enhancing pain sensitivity ^23^. However, antagonists targeting the CCL2-CCR2 axis had not generated favorable outcomes in clinical trials ^24^. HMGB1 is a member of the Damage-Associated Molecular Pattern (DAMP) family molecules ^25^. Upon inflammatory conditions, HMGB1 initiates and augments inflammatory response by recruiting inflammatory cells and producing inflammatory cytokines ^26–28^. Several different strategies targeting HMGB1 ligand and receptors have been developed and demonstrated beneficial effects in preclinical models of inflammatory diseases ^29^. Future studies should test the therapeutic potential of CCL2 and HMGB1 blockade when these inhibitors are developed to a clinical setting.

Our long-term goal is to develop effective therapy that both controls tumor growth and alleviates pain for SWN patients. We found that IL-6 neutralizing antibody reduced pain in the orthotopic PDX mouse models. IL-6 is a major trigger for the onset and progression of inflammatory diseases such as rheumatoid arthritis, inflammatory bowel disease, and sepsis^17,30,31^. The US FDA has also approved anti-IL6 (siltuximab) and anti-IL-6R antibodies (tocilizumab and sarilumab) for rheumatoid arthritis, to manage pain ^32,33^. Based on our findings, we have now initiated a clinical trial studying the activity of siltuximab in patients with SWN and moderate to severe pain (NCT05684692). Another important implication of our findings is that IL-6 may serve as clinically relevant biomarkers for pain for future clinical studies.

We identified EGF signaling as a driver of SWN progression. The ErbB family of receptor tyrosine kinases comprises the EGFR/Her1, Her2 (ErbB2), Her3 (ErbB3), and Her4 (ErbB4)^34^, which are known to contribute to the pathogenesis and progression of many human cancers ^35,36^. In patients with vestibular schwannomas, EGFR-targeted therapy has been tested. Lapatinib, an HER-1/2 inhibitor, reduced tumor size in 4 of 17 patients and had a hearing response in 4 of 13 patients ^37^. A possible reason for the limited hearing response might be that neuroinflammation, a contributing factor to hearing loss, is left unblocked. In SWNs, targetable ErbB2 mutations have been identified in SWN ^38^, however, its role in SWN progression remains to be investigated. We demonstrate here that combined IL-6 and EGFR blockade can concurrently reduce pain and inhibit tumor growth. We suggest that further clinical studies should test the combined anti-tumor and anti-inflammation strategy that can concurrently inhibit tumor growth and alleviate pain.

## Methods

Tumor-induced pain response and treatment efficacy were studied in two orthotopic PDX models. For additional information regarding cell lines, animal models, treatment protocols, patient characteristics, and statistical analysis, see Supplemental Information.

## Supplementary Materials

### Methods

Figure. S1. Characterization of pain response in male and female mice, motor function and histological confirmation of schwannoma.

Figure. S2. Peripheral nerve schwannoma triggers macrophage influx into the DRG.

Figure. S3. HMGB1 expression in SWN cells and gene knockdown effects.

Figure S4. Combined IL-6 and EGFR blockade effects on bodyweight loss and tumor growth in vivo.

Table S1. Demographics and clinical characteristics of SWN tumor used for the establishment of patient-derived SWN cell lines

Table S2. Differentially expressed genes in patient SWN tumors compared to normal Schwann cells.

Table S3. Differentially expressed genes in tumor-associated macrophage after anti-IL-6 treatment.

Table S4. Primer sequences used in qRT-PCR assay.

## Acknowledgments

We thank Mark Duquette and Anna Khachatryan for their superb technical support, Dr. Peigen Huang for assisting in animal studies, and Greg Wojtkiewicz for acquiring the MRI images.

## Author contributions

L.X., S.R.P., and J.M. designed the research; Z.Y., L.W., and Y.Z. established patient-derived cell lines; Z.Y., and Y.S. performed mouse model studies; Z.Y., J.R., G.X., and J.W.C. performed imaging studies; S.S. and W.H. analyzed RNASeq data; G.Y.L., and A.W. design and 3-D printed the DRG window; G.Y.L. created all of the schematic illustrations; N.Z., and G.B.F. established the DRG slice culture model; L.W., Y.S., and A.S.R. performed patient sample histological analysis; L.X., Z.Y., L.W., Y.Z., Y.S., J.W.C., S.S., W.H., A.M., L.Z., A.S.R. analyzed data; L.X., S.R.P., and J.M. wrote the paper. L.X. supervised the research.

## Funding

This study was supported by the NIH R01-NS126187 (to L.X. and J.M.), Department of Defense New Investigator Award (W81XWH-16-1-0219, to L.X.), Investigator-Initiated Research Award (W81XWH-20-1-0222, to L.X.), Clinical Trial Award (W81XWH2210439, to P.S.R and L.X.), Children’s Tumor Foundation Drug Discovery Initiative (to L.X.), American Cancer Society Mission Boost Award (to L.X.), and R01-NS103998 (to J.W.C.).

## Conflict of interest

None

## Notes

### Competing Interest Statement

The authors have declared no competing interest.

