## Supplemental Information for "Co-Targeting IL-6 and EGFR signaling for the treatment of schwannomatosis and associated pain"

**Supplementary Materials**

**Supplementary Materials and Methods:**

**Reagents.** Dulbecco's Mod. of Eagle's Medium and Ham's F-12 50/50 Mix were obtained from Corning (Manassas, VA). Leibovitz's L-15 Medium (L15), Neurobasal TM-A Medium (Neurobasal A), and 2% B-27™ Supplement (50X) were obtained from Thermo Fisher Scientific (Cambridge, MA). Collagen type I High Concentration, was obtained from Corning (Bedford, MA). Poly-L-lysine solution was obtained from Sigma Aldrich (Natick, MA). Schwann cell medium (SCM) and growth supplement (SCGS), were obtained from ScienCell (Carlsbad, CA).

**Patient-derived schwannomatosis cell lines.** Clinical samples and data were used in accordance with the protocol approved by the IRB at Massachusetts General Hospital (MGH). Patient samples were obtained from the Departments of Surgery and Pathology at MGH from December 2016 through July 2022. All patients gave signed informed consent for the collection of excessive tumor samples and molecular analysis. Schwannomatosis diagnosis was confirmed by Pathology. Tumor samples were collected after surgical resection and transported to the lab. In a laminar flow hood, the tissue was finely minced and centrifuged for 5 mins at 1,000 rpm. To dissociate the cells, the tissue was digested in 1 mg/mL collagenase for 3 hours at 37°C. After passing through a 70 µm strainer, the cells were collected by centrifugation for 5 mins at 1,000 rpm. The cell pellet was resuspended in an SCM medium. Isolated cells were plated on poly D lysine-coated coverslips in a 24-well plate. 3 days later, cells were immortalized with pLenti SV40 (Abm, Richmond, BC) and pLenti-EF1α-hTERT (Abm, Richmond, BC) following manufacturers' instructions [1].

*Confirmation of Schwann cell origin.* 8,000 cells were seeded in chamber slides and cultured overnight. The slides were fixed with 4% paraformaldehyde (PFA). The fixed cells were blocked with 5% goat serum and then incubated with anti-S100 and anti-Sox 10 (1:10, both are from Biocare Medical, Pacheco, CA) overnight at 4°C. Alexa 647-conjugated secondary antibodies were used for signal detection.

**Orthotopic patient-derived xenograft SWN mouse models.** All animal procedures were performed following the guidelines of the Public Health Service Policy on Humane Care of Laboratory Animals and approved by the Institutional Animal Care and Use Committee of the MGH. All patient-derived SWN cells were implanted in 8-12 weeks old immune-deficient nude mice. SWN grows on the peripheral nerves (in 89% of patients) and spine (in 74% of patients) [2]. To recapitulate SWN in the clinical setting, we have established both sciatic nerve and spine mouse models.

*Sciatic nerve model:* To mimic schwannomas grown on the peripheral nerve, A total of 3 µl of tumor cell suspension ($5x10^4$ cells/mice) was injected slowly (over 45-60 seconds) under the sciatic nerve sheath using a Hamilton syringe to prevent leakage [3,4]. Tumor formation was confirmed by 3D ultrasound imaging (Fig 1A). Sciatic nerve tumor size was measured by caliper every 3 days until tumors reached 1 cm in diameter.

*Spine model:* to mimic schwannomas grown in the spine, we injected SWN cells intrathecally ($1x10^4$ cells/mice, in 2 µl) using a 30-gauge needle between the dorsal aspects of T13 and L1. Tumor formation in the intradural and extramedullary spine space was confirmed by MR imaging and H&E staining (Fig 1D-E).

To confirm the formation of tumors in the correct anatomic location, we performed:

*Ultrasound*: To confirm the formation of sciatic nerve tumors, 3D ultrasound was performed on a Vevo 2100 Imaging System (FUJIFILM VisualSonics, Inc, Toronto, Canada) [3].

*MRI:* To confirm the formation of spine tumors, MRI of the spinal cord was performed on an animal 4.7-T MRI scanner (Bruker, Billerica, MA) with a mouse body coil under respiration-monitored isoflurane anesthesia. T1-weighted images were obtained before and after the intravenous administration of 0.3mmol/kg of Dotarem (Guerbet, Princeton, NJ) via the tail vein, using the rapid acquisition relaxation enhanced (RARE) sequence: repetition time (TR) = 700 ms, echo time (TE) = 14.0548 ms, matrix size 256 × 256, and slice thickness 0.5 mm. 12 sections were acquired [5].

**Measurement of tumor growth.** Tumor cell lines were infected with lentivirus encoding secreted Gaussia luciferase (Gluc), and the measurement of plasma Gluc was performed as previously described [6]. Briefly, 13 µl of whole blood was collected from a slight nick on tail veins and mixed with 5 µl of 50 mM EDTA immediately to avoid clotting. The blood sample was transferred to a 96-well plate, and Gluc activity was measured using a plate luminometer (GloMax 96 Microplate Luminometer, Promega). The luminometer was set to automatically inject 100 µl of 100 mM coelenterazine (CTZ, Nanolight) in PBS, and photon counts were acquired for 10 sec.

**Treatment protocols.** In the sciatic nerve model, treatment starts when the tumor reaches 3 mm in diameter. In the spine model, treatment starts when blood Gluc concentration reaches $1x10^4$ RLU. Anti-CCL2 treatment (2 mg/kg, Biolegend, San Diego, CA) was administered by *i.p.* injection, every 3 days and continued until the study endpoint. αIL-6 treatment (5 mg/kg, BioxCell, Lebanon, NH) was administered by *i.p.* injection, every 3 days and continued until the study endpoint. Dacomitinib (20 mg/kg, LC Laboratories, Woburn, MA) was administered by oral gavage 5 days/week and continued until the study endpoint.

**von Frey filament threshold testing.** von Frey filaments (DanMic Global) were used to estimate absolute withdrawal thresholds. Von Frey filaments were applied to the plantar surface of each hind paw using an ascending series. Each filament was tested 5 times in increasing order starting with the filament producing the lowest force (0.008g). The threshold force of response is defined as the first filament that evoked at least 2 withdrawals out of 5 applications [7]. All testing took place during the light cycle, no earlier than 09:00 h and no later than 16:00 h. Although data were collected on both hind paws, only data from the hind paw ipsilateral to the tumor implantation are presented, as no significant effects of sex, drug, or genotype on the contralateral paw were observed in any experiment.

**Rotarod test.** The motor function of the mice was assessed using an automated Rotarod (Columbus Instruments, Columbus, OH). Rotarod test was performed in animals bearing size-matched tumors. Tumor size was measured every 3 days by caliper; when tumor size reached approximately 6 mm in diameter, mice were trained on the rotarod every day for 3 days. Three days later, the tumor reached approximately 1 cm in diameter, and we performed rotarod test; three tests were performed for each animal: each test began with a 30-second acclimation period at 4 rpm followed by acceleration by 4 rpm every 60 seconds to a maximum of 10 min and 40 rpm. The amount of time that elapsed before the mouse fell off was recorded as rotarod endurance. To avoid heterogeneity between animals, the average time to fall from the rotating cylinder was normalized to the value from each mouse on the first day and presented as relative rotarod endurance [8].

**Adoptive transfer of bone marrow-derived macrophages.** We harvested bone marrow cells from long bones of *Ccr2*$^{-/-}$/C57BL6, *Ccr2*$^{RFP/RFP}$/C57BL6 mice and cultured with M-CSF (20 ng/ml) for 7 days to induce them into macrophages. Nude mice (8 weeks of age) were sub-lethally whole-body irradiated for a single dose of 6 Gy. Twenty-four hours after irradiation, 1x10$^6$/in 100 µl donor bone marrow-derived macrophages were delivered to the irradiated recipients through retro-orbital injection [4]. Peripheral blood was collected 3 weeks after cell injection and analyzed for the presence of RFP$^+$ and CD45.2 cells by flow cytometry analysis. Flow cytometric data were collected and analyzed using FlowJo software.

**Implantation of DRG window.** A small incision was made in the dorsal skin at the lumbar (L3-L4) level of the spine, and the skin was held back with retractors. Under a stereomicroscope, muscles, and ligaments attached to the lateral aspects of two vertebrae were detached using surgical scissors. Using a high-speed micro-drill, articular processes around DRG were gently trimmed to the same level as the DRG surface. The custom-designed, 3D-printed DRG imaging window holding a 3-mm diameter cover glass is sutured with the muscles.

**Intravital multiphoton laser scanning microscopy (MPLSM) through DRG window.** Mice bearing DRG window were anesthetized and placed on an imaging stage and scanned by a focused laser beam onto the DRG window while the reflected optical signals were recorded. To minimize breathing artifacts during imaging, the custom DRG window was fixed to a stainless-steel stage that holds the mouse in place throughout image acquisition (Fig 2B). To avoid visual artifacts induced by DRG window placement, such as fibrotic tissue accumulation from scarring, all image stacks were analyzed at a depth of 50 µm or deeper from the surface, typically between 100 and 500 µm. Intravital MPLSM imaging was performed following *i.v.* injection of 0.1 ml FITC-dextran (MW=2 million, Sigma) [9]. For each mouse, four to five cross-sectional images were provided in real-time, and three-dimensional profiles (150 µm stacks) representing tumor vessels and macrophages were obtained through the DRG window. The same regions identified by using landmarks of the tumor vasculature were recorded at different time points (Fig 2B) [9].

**HMGB1 knockout using CRISPR-Cas9 construct.** The HMGB1 CRISPR/Cas KO plasmid was purchased from Santa Cruz Biotechnology Inc (sc-400735, CA). The HMGB1 CRISPR/Cas9 KO plasmid consisted of a pool of three plasmids, each encoding the Cas9 nuclease, and the HMGB-1 specific 20-nt guide RNA (gRNA) designed for maximum KO efficiency. gRNA sequences were derived from the GeCKO (v2) library and directed the Cas9 protein to induce a site-specific double-strand break in genomic DNA. After transfection of the plasmid, single-cell clones were picked after antibiotic selection and screened for their knocked-down expression of HMGB1 by ELISA (R&D Systems). To avoid clonal variation, 3 clones with the lowest HMGB1 expression were pooled and used in the animal experiment.

**RNASeq.** For the cohort of fresh SWN samples used for bulk RNASeq analysis, surgically resected schwannoma tissue was obtained from patients with well-characterized clinical cases of SWN (Table S1). Three RNA samples from normal Schwann cells were used as a control in the RNASeq analysis. Tissues were homogenized using the polytron PT1300 tissue homogenizer, followed by additional homogenization using a Qiashredder spin column. RNA from tumor tissues was extracted using the RNeasy Mini Kit (QIAGEN, Cambridge, MA). 1 µg of total RNA was sent to Molecular Biology Core Facilities, Dana-Farber Cancer Institute. RNASeq analysis was performed following the routine procedure in the Xu lab [4]. The DESeq2 package in R was used to determine the differentially expressed genes (DEGs) [10]. To control for

False discovery rate (FDR) at 0.05, we used the Benjamini & Hochberg algorithm. ComplexHeatmap package was used to plot the heatmap [11]. The differentially expressed gene set was analyzed by Gene Set Enrichment Analysis software (GSEA, https://software.broadinstitute.org/ software/cprg/?q=node/14).

**DRG slice culture.** DRGs were surgically dissected from 1-2 day postnatal nude mice. The mice were sacrificed by decapitation. DRG was collected from L1-L6 lumbar vertebrae under the microscope and transferred to an ice-cold L15 medium. Then the DRG was cultured in a slide chamber coated with poly-L-lysine and collagen. DRG was maintained in the Neurobasal A supplemented with B27 and glutamine (both are from Thermo Fisher Scientific, Cambridge, MA), 50 ng/ml NGF (Millipore, Cambridge, MA), and 10 μM 5FDU and 1μM Cytosine α-D-arabinofuranoside (AraC, Sigma Aldrich, Natick, MA). To collect tumor cell-conditioned medium from tumor cells or macrophages, cells were grown in one T75 flask at 50% confluency in 10 ml growth medium. 3 days later, the medium was harvested and centrifuged at 1,500 g and sterile filtered (0.22 μM filter).

**Isolation of TAMs and peritoneal macrophages**. Tumor-associated macrophages were isolated by fluorescence-activated cell sorting. Briefly, tumor tissues were mechanically chopped and enzymatically digested for 1 hour at 37°C with 3mg/ml collagenase A (Roche) in serum-free DMEM medium. Digestion was stopped by the addition of DMEM supplemented with 8% FBS and the suspension was disaggregated through a 70 μm cell strainer. Single-cell suspension was stained for 20 min at 4°C with anti-mouse F4/80 (1:200, BM8; eBioscience). Sorting of F4/80$^{+}$ cells was performed on a BD FACS sorter. Peritoneal macrophages were collected by peritoneal lavage from mice given an intraperitoneal injection of 2 ml of thioglycolate broth (Sigma) following our previously published protocol [12]. Briefly, 4 days after thioglycolate injection, peritoneal macrophages were harvested and washed with $Ca^{2+}$- and $Mg^{2+}$-free PBS, and suspended in 10% DMEM for culture and in vitro treatment [12].

**Gene expression analysis.**

*Quantitative RT-PCR.* qPCR was performed to evaluate changes in mRNA level using SYBR Green-based protocol [13]. All qPCR and analysis were performed on a Stratagene MX 3000 qPCR System operating MXPro qPCR software (Stratagene, San Diego, CA) [14].

*Western Blot.* Thirty micrograms of protein per sample were separated on 10% SDS-polyacrylamide gels [15]. Membranes were blotted with antibodies against total (1:500) and phospho-ErbB2 (1:1000); total (1:500) and phospho-ErbB3 (1:500); total (1:500) and phospho-ErbB4 (1:500); total (1:1000) and phospho-Akt (1:200); total and phospho-S6 (1:1000 for both). Antibodies were obtained from Cell Signaling (Danvers, MA). Membranes were blotted with beta-actin for equal loading control (1:5000, Sigma) [16].

*ELISA.* Plasma or protein extracted from snap-frozen tumors were diluted to 2 μg/μl concentration according to protein assay. Mouse inflammatory cytokine levels were quantified using mouse multiplex enzyme-linked immunosorbent assay plates following the manufacturer's instructions (Meso-Scale Discovery, Gaithersburg, MD). Every sample was run in triplicate [9,13].

**MTT assay.** Five thousand cells were seeded into 38 $mm^2$ wells of flat bottomed 96-well plates in triplicate and allowed to adhere overnight. 3-(4,5-Dimethylthiazol-2-yl)-2,5-diphenyltetrazolium bromide (MTT, 5 mg/mL, Sigma Chemical) was prepared in PBS. The number of metabolically active cells was determined by MTT assay [17].

**Histological staining.** DRG slices were fixed with 4% PFA and 20% sucrose, blocked in 5% BSA for 1 hour, and then incubated against anti-CGRP (1:5000) and anti-TRPV1 (1:800, both are from Abcam, Cambridge, MA)[9]. To evaluate tumor cell proliferation and apoptosis, slides of tumor tissues were stained with proliferating cell nuclear antigen (PCNA, 1:1000, Abcam) and TUNEL (ApopTag® Peroxidase *In Situ* Apoptosis Detection Kit, EMD Millipore) following the manufacturer's instructions. Positively stained cells were manually counted [18]. The infiltration tumor-associated macrophages were stained by Iba1 (1:5000, Wako). Tumor-infiltrating macrophages and expression of IL-6 pathway genes were evaluated by immunofluorescent staining using archived paraffin-embedded Schwannomatosis patient samples with antibodies against CD163 (PGM1; Dako, Santa Clara, MA), IL-6 (1:100, Abcam, Cambridge, MA), and phosphor-Stat3 (1:10, Cell Signaling Technology, Danvers, MA). Four specimens of normal peripheral nerve were obtained postmortem and used as controls. Slides were incubated overnight at $4^{o}$C. Sections are counterstained with DAPI and mounted. Appropriate positive and negative controls were used for all stains.

*Image quantification:* For the macrophage, PCNA, and TUNEL staining, positively stained cells are manually counted. The positively stained area of all other histological staining was evaluated with digital quantitative image analysis using the open-source software ImageJ. Positive staining in 20 random fields/slides was quantified via automated built-in functions based on fluorescent pixel intensity after establishing a threshold to exclude background staining. Individual staining was quantified as area fractions of the tumor region of interest and reported as percentages [4].

**Statistical analyses.** Spearman's rho correlation coefficient was calculated. We determined whether growth curves significantly differed from each other by log-transforming the data, fitting a linear regression to each growth curve, and comparing the slopes of the regression lines (using an equivalent of ANOVA). Significant differences between the two groups were analyzed using the Student's t-test (two-tailed) or Mann-Whitney *U* test (two-tailed). All calculations were done using GraphPad Prism Software 6.0 and Microsoft Excel Software 2010. Differences in hearing response between multiple groups were evaluated with ANOVAs. Welch's t-test was used for comparison between the two groups. Benjamini-Hochberg correction for multiple comparisons was applied. Histological analysis of gene expression in patient schwannoma tissues was statistically analyzed using statistics software (Statistical Package for the Social Sciences Statistics, Version 20.0; IBM) with an a priori significant P value < 0.05.

### Supplementary Figures

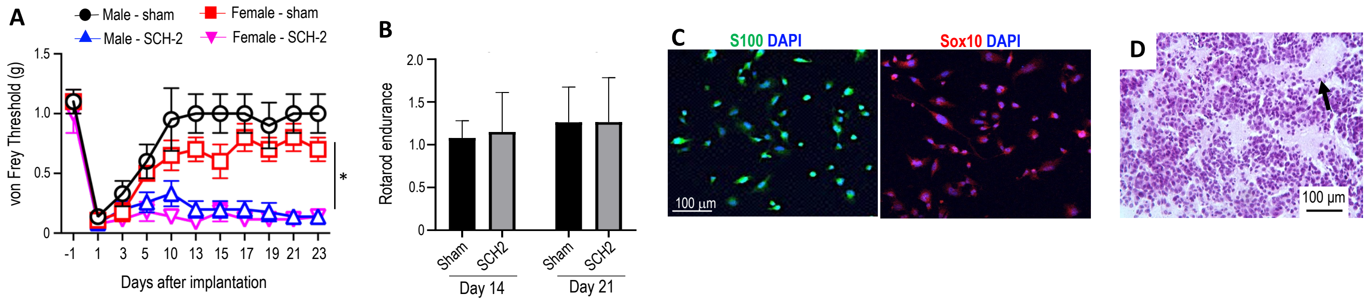


**Figure S1. Characterization of pain response in male and female mice, motor function and histological confirmation of schwannoma. (A)** von Frey filament test was performed in male or female mice receiving sham surgery or bearing SCH-2 tumors in the sciatic nerve. N=12/ea, *P<0.001. **(B)** Rotarod test was performed in SCH-2 sciatic nerve model on Day 14 and 21 post-tumor implantations. N=12/ea. **(C)** Representative IF staining images of Schwann cell markers S100 (green) and Sox 10 (red), DAPI (blue). **(D)** Representative H&E staining of sciatic nerve SCH-2 tumors demonstrating classic schwannoma histology features of Verocay body (arrow pointed).

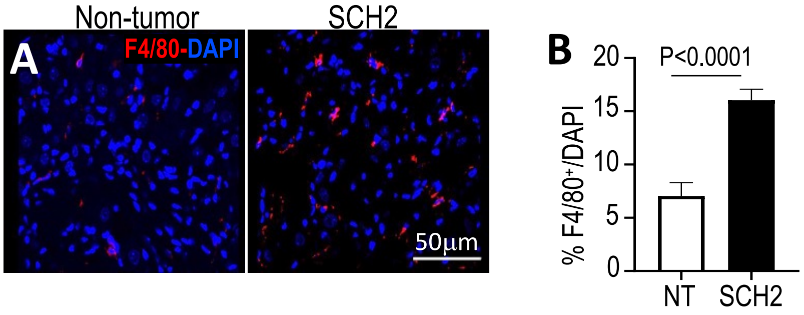


**Figure S2. Peripheral nerve schwannoma triggers macrophage influx into the DRG**. **(A)** Representative images of IF staining for macrophages (F4/80, red) in the DRGs from non-tumor bearing mice and SCH2 tumor-bearing mice. DAPI, blue. **(B)** The percent of positively stained cells was manually counted. Non-tumor bearing, n=8 mice, SCH-2, n=14 mice. 20 random areas were imaged and quantified. Data presented as mean±SD.

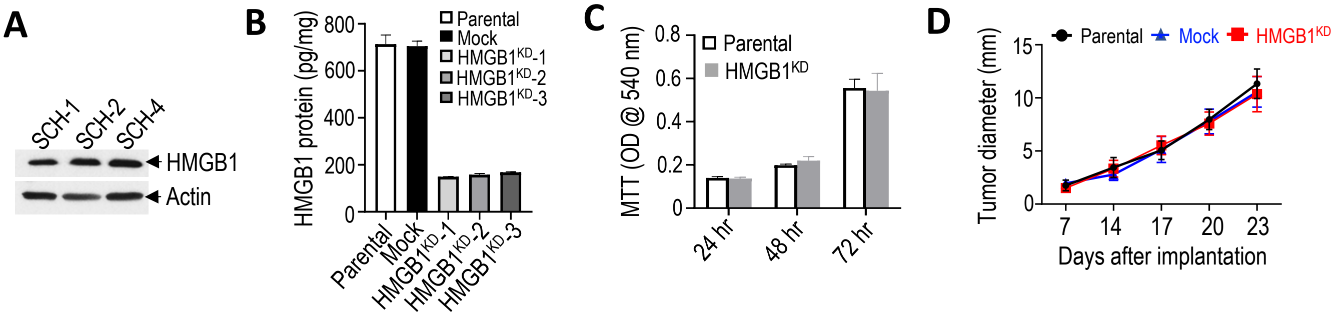


**Figure S3. HMGB1 expression in SWN cells and gene knockdown effects. (A)** Western blot analysis of HMGB1 protein in patient-derived schwannoma cells *in vitro*. Comparison between parental, mock-transfected, and 3 clones of HMGB1 knockdown cells in their **(B)** HMGB1 protein level by ELISA, and **(C)** in vitro cell viability by MTT assay. **(D)** Tumor diameter measured by caliper. N=8 mouse/group. Animal studies are presented as mean±SEM, and are representative of at least three independent experiments.

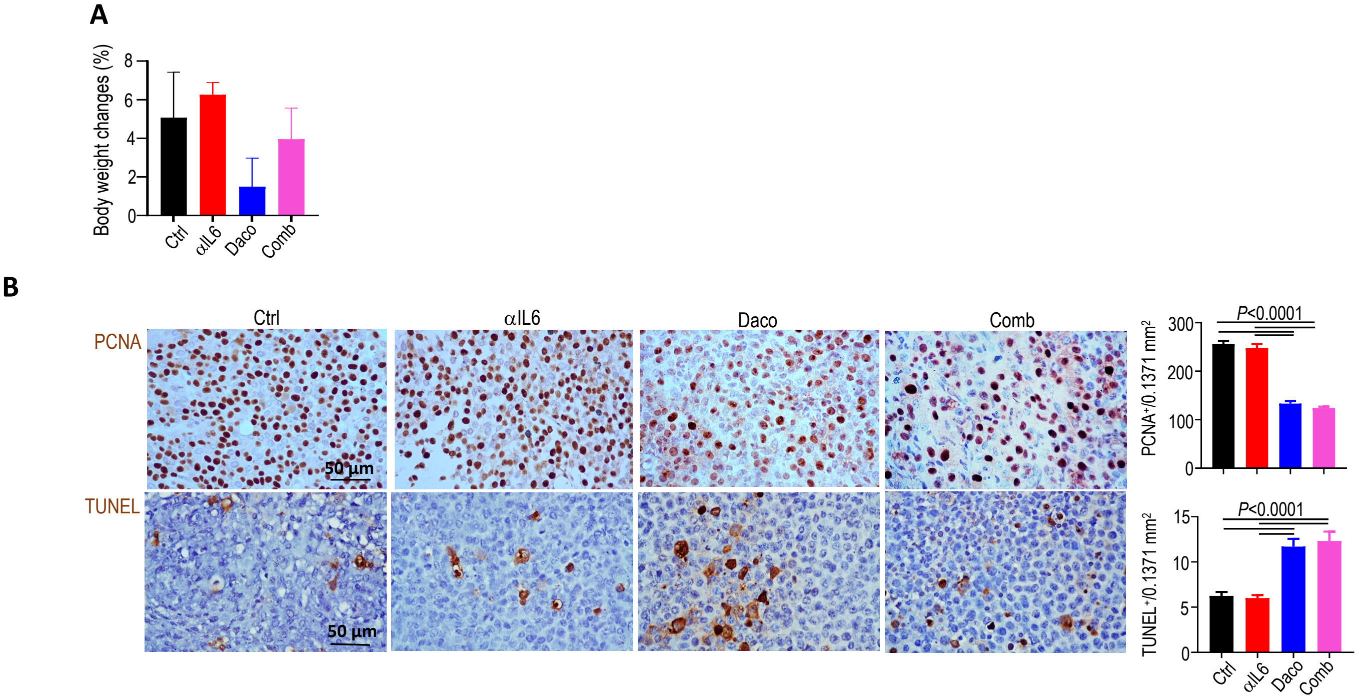


**Figure S4. Combined IL-6 and EGFR blockade effects on bodyweight loss and tumor growth in vivo. (A)** Body weight is monitored every 3 days, and data collected at the experiment endpoint is presented. N=12/group. **(B)** Representative IHC images of PCNA to mark proliferating cells and TUNEL to mark apoptotic cells. **(B)** Quantification of IHC staining. N=8 mice/group, 20 random areas were imaged and quantified. Data present mean ± SD.

**Table S1. Demographics and clinical characteristics of SWN tumor used for the establishment of patient-derived SWN cell lines**

| Cells | Sex | Age | Pain status | Tumor location | Tumor size at surgery (mm) |
|---|---|---|---|---|---|
| **SCH-1** | M | 36 | Non-painful | Left accessory nerve/schwannoma | 26 |
| **SCH-2** | F | 39 | Painful | Right pudendal nerve/schwannoma | 37 |
| **SCH-4** | M | 60 | Painful | Left tibial nerve/schwannoma | 45 |
| **SCH-5** | F | 57 | Painful | Right thigh perineuroma | 34 |
| **SCH-6** | F | 41 | Painful | Left common peroneal nerve/schwannoma | 59 |
| **SCH-7** | F | 40 | Non-painful | T5 meningioma | 15 |
| **SCH-9** | M | 38 | Non-painful | T12 Schwannoma | 19 |
| **SCH-10** | M | 53 | Non-painful | Presacral/schwannoma | 66 |
| **SCH-11** | F | 49 | Painful | Cervical/schwannoma | 29 |

**Table S2.** **Differentially expressed genes in patient schwannomas compared to normal Schwann cells**

| Gene_name | logFC | PValue | FDR |
|---|---|---|---|
| DDX3Y | -10.843099 | 9.59E-17 | 1.09E-12 |
| CCL13 | -14.104544 | 1.39E-12 | 7.90E-09 |
| CCL11 | -14.207724 | 2.92E-09 | 1.10E-05 |
| MMP9 | -7.4621405 | 4.01E-09 | 1.14E-05 |
| XIST | 8.76997557 | 7.28E-09 | 1.66E-05 |
| MFAP5 | -8.4085353 | 2.13E-08 | 3.48E-05 |
| CXCL8 | -5.5695175 | 2.15E-08 | 3.48E-05 |
| NKX2-1 | 12.2419577 | 3.66E-08 | 4.62E-05 |
| CHI3L1 | -10.443753 | 3.66E-08 | 4.62E-05 |
| ACAN | -7.2772828 | 8.19E-08 | 9.31E-05 |
| KDM5D | -7.1657084 | 1.56E-07 | 0.00016135 |
| TRH | -16.171674 | 1.70E-07 | 0.00016135 |
| EFEMP1 | -6.4902765 | 2.52E-07 | 0.00021579 |
| FMOD | -6.3723571 | 2.66E-07 | 0.00021579 |
| MEDAG | -6.7746594 | 5.49E-07 | 0.00041598 |
| HSD11B1 | -8.2399562 | 6.24E-07 | 0.00043032 |
| S100A1 | -6.3589894 | 6.43E-07 | 0.00043032 |
| AKR1C2 | -6.9894431 | 6.89E-07 | 0.00043533 |
| HAND2 | -9.3627706 | 7.75E-07 | 0.00044685 |
| ADRA2C | -8.1520488 | 7.86E-07 | 0.00044685 |
| LOC1019288 | 4.81293458 | 1.40E-06 | 0.00070821 |
| CXCL6 | -10.403654 | 1.40E-06 | 0.00070821 |
| ACTA2 | -5.7040494 | 1.43E-06 | 0.00070821 |
| OSR2 | -5.3853529 | 1.93E-06 | 0.00091506 |
| PRG4 | -8.0996517 | 2.11E-06 | 0.00095927 |
| COMP | -8.7640345 | 2.39E-06 | 0.0010443 |
| LINC01133 | -12.597176 | 2.60E-06 | 0.00109646 |
| ELN | -7.4206491 | 3.05E-06 | 0.00123665 |
| CRYBA2 | -10.435479 | 3.47E-06 | 0.00136141 |
| NDUFA4L2 | -5.2294122 | 3.62E-06 | 0.00136369 |
| ITGBL1 | -3.6155406 | 3.72E-06 | 0.00136369 |
| AKR1C1 | -5.6687608 | 3.87E-06 | 0.00137374 |
| IFI27 | -4.7874654 | 4.12E-06 | 0.0014197 |
| ABI3BP | -4.2549294 | 4.40E-06 | 0.00147174 |
| CH25H | -4.3563198 | 4.92E-06 | 0.00159683 |
| SVEP1 | -4.1390312 | 5.46E-06 | 0.00167128 |
| PTGS2 | -4.1234695 | 5.54E-06 | 0.00167128 |
| RPS4Y1 | -7.8052783 | 5.59E-06 | 0.00167128 |
| CXCL1 | -7.5741634 | 6.26E-06 | 0.00182526 |
| SCG2 | -5.157 | 8.05E-06 | 0.00228883 |
| BARHL2 | 11.1793231 | 9.26E-06 | 0.00256692 |
| PID1 | -5.3255509 | 9.98E-06 | 0.0027026 |

| | | | |
|---|---|---|---|
| PDGFRA | -4.6499168 | 1.05E-05 | 0.0027365 |
| AKR1C3 | -6.5623886 | 1.06E-05 | 0.0027365 |
| IFNG | 5.09616408 | 1.12E-05 | 0.00283846 |
| CRABP2 | -5.0213142 | 1.20E-05 | 0.00295774 |
| PAMR1 | -4.7121742 | 1.54E-05 | 0.003646 |
| IL6 | 5.73838057 | 1.54E-05 | 0.003646 |
| BMP6 | -3.5313717 | 1.64E-05 | 0.00380162 |
| MIR6723 | 6.17674328 | 1.91E-05 | 0.00430864 |
| LOC1019304 | -6.2575984 | 1.93E-05 | 0.00430864 |
| TNNT2 | -7.7980123 | 2.08E-05 | 0.00451173 |
| ABLIM1 | -3.3775179 | 2.10E-05 | 0.00451173 |
| TAGLN | -5.3380574 | 2.25E-05 | 0.00472994 |
| ANGPTL7 | -4.7555953 | 2.47E-05 | 0.0051029 |
| LXN | -4.1871908 | 2.68E-05 | 0.00544626 |
| SRPX2 | -3.3767159 | 2.76E-05 | 0.0054953 |
| CCL7 | -8.1720711 | 2.90E-05 | 0.0055954 |
| CFB | -4.1094108 | 2.94E-05 | 0.0055954 |
| A4GALT | -3.7893206 | 2.95E-05 | 0.0055954 |
| MGP | -4.2806493 | 3.02E-05 | 0.0056282 |
| HMGB1 | 9.05963919 | 3.09E-05 | 0.0056723 |
| HOXB9 | 6.60902972 | 3.31E-05 | 0.00597786 |
| CCL2 | -4.8854765 | 3.86E-05 | 0.00684921 |
| MDGA1 | -4.5064069 | 3.97E-05 | 0.00694156 |
| ACTG2 | -5.2091908 | 4.14E-05 | 0.007139 |
| CDR1 | 5.70378305 | 4.30E-05 | 0.00729992 |
| PCSK1 | -5.7967148 | 5.21E-05 | 0.0087057 |
| CYP1B1 | -3.6304992 | 5.29E-05 | 0.00871369 |
| HSPB7 | -5.0165277 | 5.39E-05 | 0.00871369 |
| IGFBP3 | -4.7965356 | 5.46E-05 | 0.00871369 |
| PI16 | -6.3344684 | 5.52E-05 | 0.00871369 |
| FAM43A | -4.1541063 | 6.05E-05 | 0.00941571 |
| EN1 | -4.7700319 | 6.22E-05 | 0.00956208 |
| TXLNGY | -6.3386096 | 6.58E-05 | 0.00998112 |
| PTGES | -4.6574936 | 6.77E-05 | 0.01013087 |
| MAB21L1 | -4.498286 | 7.08E-05 | 0.01045603 |
| DBNDD1 | -3.4644042 | 7.98E-05 | 0.01162482 |
| PTPRN | -6.6087502 | 9.05E-05 | 0.01302164 |
| USP9Y | -6.5009366 | 9.37E-05 | 0.01331423 |
| PRRX2 | -6.029969 | 0.00010649 | 0.01494723 |
| PDE1A | -4.7204378 | 0.00010985 | 0.01512455 |
| OLFML1 | -4.7264367 | 0.00011042 | 0.01512455 |
| RARRES1 | -4.0504325 | 0.00011454 | 0.01550304 |
| NSG1 | -4.239188 | 0.00011764 | 0.01573523 |
| TP53I11 | -4.4982026 | 0.00012248 | 0.01602405 |
| TRPV4 | -5.6543735 | 0.00012262 | 0.01602405 |

| | | | |
|---|---|---|---|
| ZNF423 | -4.6130339 | 0.00012595 | 0.0162691 |
| CSF3 | -8.6094622 | 0.00012736 | 0.0162691 |
| FAM20A | -4.5493815 | 0.00013109 | 0.01637314 |
| PTGIS | -6.1327566 | 0.00013353 | 0.01637314 |
| SPEG | -4.4137915 | 0.00013355 | 0.01637314 |
| PTHLH | -5.9733668 | 0.00013393 | 0.01637314 |
| HYAL1 | -4.1717036 | 0.00014182 | 0.01715219 |
| SCRG1 | -5.1489505 | 0.00014569 | 0.01739155 |
| CNN1 | -4.2106587 | 0.00014685 | 0.01739155 |
| LRAT | -4.4832178 | 0.00014871 | 0.01742946 |
| KLHDC8A | 6.29293802 | 0.000151 | 0.01751807 |
| COL4A4 | -4.495426 | 0.00016192 | 0.01859458 |
| ISLR | -3.4153945 | 0.00017417 | 0.01980186 |
| NOV | -4.4946312 | 0.00018164 | 0.02044596 |
| PDGFRL | -3.6380708 | 0.00019841 | 0.02211512 |
| BDKRB2 | -5.7428576 | 0.00020333 | 0.02244345 |
| MEOX2 | -4.1805248 | 0.00021558 | 0.02356691 |
| SULF1 | -4.5359883 | 0.00022133 | 0.0239645 |
| CD70 | -6.9592043 | 0.00022572 | 0.02420939 |
| H19 | -3.8023272 | 0.00023219 | 0.0246709 |
| CAMK2B | 4.59672741 | 0.00023842 | 0.02509864 |
| LIF | -4.0721775 | 0.00024995 | 0.02607089 |
| NXPH4 | -6.1045505 | 0.00026241 | 0.02712098 |
| ARHGEF16 | 4.15768222 | 0.00026768 | 0.02741704 |
| EFNB2 | 2.61266057 | 0.00027356 | 0.02760783 |
| TGFBI | -4.6928485 | 0.0002744 | 0.02760783 |
| CNTN1 | -5.0435706 | 0.00027703 | 0.02762814 |
| DIAPH3 | 3.28567863 | 0.0002815 | 0.02782906 |
| IL20RB | -4.2813487 | 0.00028631 | 0.02790988 |
| GPC6 | -3.1471601 | 0.00028722 | 0.02790988 |
| CGNL1 | -4.3858537 | 0.0002926 | 0.02819114 |
| SNHG5 | -2.6876028 | 0.00029691 | 0.02836584 |
| CST1 | -6.6171252 | 0.00030173 | 0.02858644 |
| RAP1GAP | 4.73118893 | 0.00031002 | 0.02912949 |
| HAPLN2 | 6.81831565 | 0.00032366 | 0.03006514 |
| EIF1AY | -5.3230347 | 0.00032527 | 0.03006514 |
| CHODL | 6.59124778 | 0.0003372 | 0.0308986 |
| GALNT12 | -4.5052665 | 0.00033972 | 0.0308986 |
| MMP3 | -9.0675199 | 0.00038224 | 0.03443782 |
| MRGPRF | -3.1194674 | 0.00038671 | 0.03443782 |
| ADGRB1 | 5.89801464 | 0.00038772 | 0.03443782 |
| NPAS1 | -4.7141542 | 0.00039556 | 0.03455779 |
| CDH7 | 6.4408788 | 0.0003987 | 0.03455779 |
| SLIT3 | -3.9944992 | 0.00039976 | 0.03455779 |
| MEGF6 | -2.6989105 | 0.00040717 | 0.03455779 |

| | | | |
|---|---|---|---|
| ADRB2 | 3.90649966 | 0.00040797 | 0.03455779 |
| MSC | -4.3445856 | 0.00040906 | 0.03455779 |
| TIMP1 | -3.6907713 | 0.00041035 | 0.03455779 |
| IGFBP5 | -3.7756272 | 0.00042475 | 0.03550732 |
| ABCB5 | -6.0191102 | 0.000432 | 0.03584982 |
| IGFBP7 | -3.0532938 | 0.00044374 | 0.03655721 |
| DRP2 | 4.73665309 | 0.00045562 | 0.03726575 |
| TNXB | -4.6074754 | 0.00046065 | 0.03737441 |
| SNED1 | -3.1199928 | 0.00046352 | 0.03737441 |
| RGS16 | -3.2708267 | 0.00047361 | 0.03791886 |
| COL22A1 | -4.3408083 | 0.00048389 | 0.03838128 |
| CRLF1 | -4.1730517 | 0.00048614 | 0.03838128 |
| XPNPEP2 | -5.2227811 | 0.00048954 | 0.03838353 |
| PKDCC | -3.6295489 | 0.00050045 | 0.03892929 |
| SFRP2 | -4.9489294 | 0.00050335 | 0.03892929 |
| GNA14 | -3.898569 | 0.00051585 | 0.03962611 |
| IL1A | -5.2183617 | 0.00052245 | 0.03986361 |
| KRT18 | -5.5282679 | 0.00054145 | 0.04103809 |
| CXCL12 | -4.16205 | 0.00056004 | 0.04216641 |
| NUPR1 | -3.4586683 | 0.00057259 | 0.04227876 |
| MASP1 | -6.9490287 | 0.00057285 | 0.04227876 |
| MFI2 | -3.5119785 | 0.00058045 | 0.04227876 |
| ADH1B | -4.3206783 | 0.00058101 | 0.04227876 |
| UCN2 | -4.4657938 | 0.00058149 | 0.04227876 |
| HIC1 | -3.188101 | 0.00058385 | 0.04227876 |
| COL11A2 | -4.0258071 | 0.00059533 | 0.04283768 |
| EBF2 | -3.3031582 | 0.00062405 | 0.04462178 |
| LAMC2 | -4.0650568 | 0.00065659 | 0.04620917 |
| SHOX2 | -3.9505788 | 0.00065695 | 0.04620917 |
| ITGA4 | 2.92820008 | 0.00066175 | 0.04620917 |
| POU3F1 | 4.84448794 | 0.00066251 | 0.04620917 |
| KCNJ10 | 6.02837232 | 0.00066809 | 0.04631411 |
| CEMIP | -4.4010612 | 0.00070071 | 0.04828108 |
| ADGRB3 | 4.7095355 | 0.00071404 | 0.04890301 |
| FLRT2 | -3.1148559 | 0.00073621 | 0.04988785 |
| ASS1 | -3.4513054 | 0.00073719 | 0.04988785 |
| ZNF185 | 3.54771443 | 0.00074531 | 0.05013894 |
| P4HA3 | -4.3622281 | 0.00076006 | 0.05083012 |
| SMAD6 | -3.2364178 | 0.0008439 | 0.05598802 |
| BDKRB1 | -6.7158743 | 0.00084703 | 0.05598802 |
| NTNG1 | -2.9249755 | 0.00089239 | 0.05864488 |
| TRHDE | 4.44149758 | 0.0008989 | 0.05873324 |
| BMPER | -5.5162092 | 0.00093386 | 0.06066867 |
| LINC01279 | -4.0408843 | 0.00094103 | 0.06078725 |
| DGKB | 3.75725346 | 0.0009504 | 0.06100627 |

| | | | |
|---|---|---|---|
| GAP43 | -4.2452688 | 0.00095515 | 0.06100627 |
| BGN | -3.7008968 | 0.00096404 | 0.06123024 |
| ADD2 | 4.36461465 | 0.00100207 | 0.06329189 |
| SPON1 | -3.1445512 | 0.00100922 | 0.06339141 |
| OLFM2 | -4.1631974 | 0.00103851 | 0.06487255 |
| FOXF1 | 5.50262847 | 0.00108663 | 0.06750773 |
| SELENBP1 | -2.9187327 | 0.00113187 | 0.06993581 |
| SOD3 | -3.2811895 | 0.00113816 | 0.06994474 |
| TNFSF9 | -3.3933432 | 0.00125954 | 0.07698748 |
| GSTM1 | -5.9030656 | 0.00129625 | 0.07880783 |
| NEFM | -3.1314339 | 0.00131982 | 0.07981396 |
| SFTA3 | 6.36215044 | 0.0013286 | 0.07991976 |
| PTX3 | -3.6248125 | 0.00136595 | 0.08173413 |
| BIRC7 | 5.0652069 | 0.00143639 | 0.08549932 |
| CDH3 | -4.1714706 | 0.0014866 | 0.08802714 |
| ADAMTS16 | 4.9586318 | 0.00150167 | 0.08845832 |
| LY6E | -2.652933 | 0.00151306 | 0.08855172 |
| FAM65B | -4.1330453 | 0.00151883 | 0.08855172 |
| HMGA2 | 4.53152947 | 0.00154979 | 0.089896 |
| SEMA3G | 4.64910366 | 0.00156233 | 0.0901632 |
| COL3A1 | -3.6414313 | 0.00158472 | 0.09099341 |
| IFITM1 | -2.6720923 | 0.00161367 | 0.09219001 |
| ADGRG1 | -2.9275997 | 0.00164745 | 0.09364934 |
| SLC9A3 | 3.54906615 | 0.00181179 | 0.10247882 |
| C11orf96 | -2.7827867 | 0.00188513 | 0.10541159 |
| SELM | -2.5124522 | 0.00189344 | 0.10541159 |
| WNK2 | 4.89233971 | 0.00190042 | 0.10541159 |
| LRRC15 | -3.235537 | 0.00190435 | 0.10541159 |
| ADGRL2 | -2.5033324 | 0.00191 | 0.10541159 |
| METTL7B | 3.86530083 | 0.00193072 | 0.10575049 |
| FOXC1 | -2.8990652 | 0.00193474 | 0.10575049 |
| FZD1 | -2.5569011 | 0.00197927 | 0.1074515 |
| GRIK2 | -3.3291274 | 0.0019999 | 0.1074515 |
| AIF1L | 3.2743209 | 0.00200105 | 0.1074515 |
| FBXO32 | -2.2130501 | 0.00200602 | 0.1074515 |
| KCNE4 | -2.8628428 | 0.00201312 | 0.1074515 |
| MFAP4 | -3.0787549 | 0.00210393 | 0.11098217 |
| COX7A1 | -3.2089894 | 0.00210489 | 0.11098217 |
| CXCL2 | -3.3154207 | 0.00211274 | 0.11098217 |
| CD248 | -3.9731027 | 0.00211832 | 0.11098217 |
| CERCAM | -2.7743986 | 0.00217119 | 0.1132305 |
| TNFRSF11B | -3.101924 | 0.00219696 | 0.1137589 |
| IGFBP4 | -2.8492502 | 0.00220133 | 0.1137589 |
| ATP13A4 | 5.70568649 | 0.00222781 | 0.1145206 |
| ATF3 | -2.4489969 | 0.00223707 | 0.1145206 |

| | | | |
|---|---|---|---|
| PDZRN3 | -2.48978 | 0.00224629 | 0.1145206 |
| HS6ST2 | 3.93819147 | 0.00228521 | 0.1159844 |
| BHMT2 | -3.12254 | 0.00231319 | 0.11688275 |
| CDH13 | 2.47205707 | 0.00233596 | 0.11751127 |
| NOTCH3 | -2.796509 | 0.00237316 | 0.11840159 |
| ARHGAP19 | 2.68868949 | 0.00237449 | 0.11840159 |
| TP53I3 | -2.339349 | 0.00245948 | 0.12210425 |
| LCTL | -3.5854531 | 0.00249169 | 0.12316557 |
| GATA6 | -3.6541827 | 0.00253124 | 0.12404636 |
| PGF | -2.9056986 | 0.00253134 | 0.12404636 |
| PLA2G4A | -2.3069926 | 0.00262121 | 0.12789938 |
| DKK1 | 2.96910371 | 0.00273499 | 0.13286658 |
| ID3 | -2.7872899 | 0.00274638 | 0.13286658 |
| SLC6A6 | -2.6496284 | 0.00296544 | 0.14285634 |
| MIR210HG | -3.6494386 | 0.00298648 | 0.14326266 |
| SLC22A3 | -4.83411 | 0.00311394 | 0.14874936 |
| BAMBI | -2.5870624 | 0.00320322 | 0.15237424 |
| GJC3 | 4.68914812 | 0.00332505 | 0.1566333 |
| RPS26 | -2.5691188 | 0.00333725 | 0.1566333 |
| PDPN | -2.6345511 | 0.00335225 | 0.1566333 |
| PODN | -3.3143172 | 0.0033541 | 0.1566333 |
| SCARA5 | -4.0020287 | 0.00336383 | 0.1566333 |
| RAP1GAP2 | 2.36284096 | 0.00337542 | 0.1566333 |
| LRRC32 | -2.4427233 | 0.00363016 | 0.16723993 |
| CHPF | -2.2800372 | 0.00363341 | 0.16723993 |
| ANKRD37 | -4.1121764 | 0.00365432 | 0.16752405 |
| PRX | 4.40909592 | 0.00367457 | 0.16777595 |
| ICAM1 | -2.2265503 | 0.00386049 | 0.17487464 |
| ARSI | -2.4677631 | 0.00386081 | 0.17487464 |
| IGFBP6 | -2.5263345 | 0.00389681 | 0.17513051 |
| LTBP2 | -2.7128353 | 0.00389727 | 0.17513051 |
| ADAMTS15 | -2.9245172 | 0.00394313 | 0.17649408 |
| EBF1 | -3.1430938 | 0.00402206 | 0.17932079 |
| ARSA | -2.1613875 | 0.004051 | 0.17990536 |
| CFH | -2.3714365 | 0.00407394 | 0.18022035 |
| FHL1 | -2.6823093 | 0.00409994 | 0.18066761 |
| CACNA1H | -3.6966584 | 0.0041255 | 0.18109179 |
| PRSS12 | -3.3737189 | 0.00424338 | 0.18503956 |
| SV2C | 5.41821244 | 0.00424798 | 0.18503956 |
| CST2 | -3.9577738 | 0.00427233 | 0.1853899 |
| MME | 2.82684467 | 0.0042897 | 0.18543555 |
| ADAM23 | -3.8708859 | 0.00431822 | 0.18596144 |
| SAMD11 | -3.5182125 | 0.00460034 | 0.19736328 |
| PFKFB4 | -2.590678 | 0.00472501 | 0.20194974 |
| CD274 | 2.94210036 | 0.00496004 | 0.21071743 |

| | | | |
|---|---|---|---|
| MRC2 | -2.7086309 | 0.00497016 | 0.21071743 |
| F3 | -2.0696073 | 0.00498575 | 0.21071743 |
| S100A13 | -2.2916569 | 0.00504001 | 0.21222171 |
| HEPH | -2.6575822 | 0.00506564 | 0.21251375 |
| PDLIM2 | -1.9975266 | 0.00508448 | 0.21252017 |
| BST1 | -2.5472617 | 0.00516985 | 0.21529663 |
| SOD2 | -2.1488232 | 0.00524649 | 0.21769094 |
| GLDC | 2.48610273 | 0.00531643 | 0.21979091 |
| SNAI1 | -2.7846708 | 0.00540631 | 0.22202841 |
| TRIM17 | 3.24870755 | 0.00540961 | 0.22202841 |
| CRTAC1 | 4.32761555 | 0.00545148 | 0.22273 |
| FBLN1 | -3.3632209 | 0.00546669 | 0.22273 |
| SLC7A8 | -3.4384667 | 0.00548548 | 0.22273 |
| TRAF1 | -2.1978471 | 0.00550573 | 0.22275686 |
| WNT2B | 2.98369118 | 0.00553797 | 0.22326654 |
| WISP2 | -3.5206812 | 0.00557421 | 0.22393335 |
| CYB5R3 | -2.2724905 | 0.005617 | 0.22485796 |
| KRT8 | -3.6338904 | 0.0057058 | 0.22648405 |
| PCOLCE | -2.9524727 | 0.00570912 | 0.22648405 |
| NRP2 | -2.2585881 | 0.00571738 | 0.22648405 |
| LOXL2 | -2.7381128 | 0.00577342 | 0.22790968 |
| SFTA1P | -5.1706355 | 0.00592623 | 0.23241487 |
| TPD52L1 | -3.1558647 | 0.00592843 | 0.23241487 |
| FOXD2 | -2.9767301 | 0.00605614 | 0.23660567 |
| PLEKHA6 | -3.0163557 | 0.00616437 | 0.24000945 |
| PLEKHA7 | 2.85909236 | 0.00633525 | 0.24582056 |
| ID1 | -3.2637498 | 0.00636421 | 0.24606319 |
| MAGI1 | 2.02235653 | 0.0064016 | 0.24606319 |
| LSAMP | -3.1529932 | 0.00641708 | 0.24606319 |
| CX3CR1 | 4.80089002 | 0.00642807 | 0.24606319 |
| CXCR4 | 2.01440769 | 0.00647463 | 0.24701367 |
| PRICKLE1 | -2.5443026 | 0.00652133 | 0.24796321 |
| PIM1 | -2.1618928 | 0.0066172 | 0.25076988 |
| ZFX | 1.94105403 | 0.0066685 | 0.25187445 |
| TMEM2 | 1.92983666 | 0.00682169 | 0.2562813 |
| TENM2 | -3.0615992 | 0.00683026 | 0.2562813 |
| NFATC4 | -2.1897585 | 0.00686289 | 0.25665865 |
| CP | -3.6515146 | 0.00698639 | 0.26042062 |
| ASIC1 | -2.6974087 | 0.00706899 | 0.26183392 |
| APLN | -2.9304761 | 0.00707037 | 0.26183392 |
| NTRK3 | 4.12235927 | 0.00712545 | 0.26285974 |
| DCN | -2.7821079 | 0.00714431 | 0.26285974 |
| NTN4 | -3.1303727 | 0.00718244 | 0.26341011 |
| GFRA1 | 1.94194255 | 0.00725422 | 0.26498163 |
| GFRA2 | 2.69675531 | 0.00730369 | 0.26498163 |

| | | | |
|---|---|---|---|
| MAP1A | -2.1011775 | 0.00731784 | 0.26498163 |
| KDELR3 | -2.242218 | 0.00736189 | 0.26498163 |
| TINAGL1 | -2.442428 | 0.00736194 | 0.26498163 |
| PDGFRB | -2.4966645 | 0.00736513 | 0.26498163 |
| NAV2 | 2.10330072 | 0.00774196 | 0.27766046 |
| PLD6 | 2.42356485 | 0.00783993 | 0.28028974 |
| GXYLT2 | -2.6160514 | 0.00788844 | 0.28114004 |
| IMPA2 | -2.4416409 | 0.00791383 | 0.28116367 |
| PAX8-AS1 | -3.1216563 | 0.00802063 | 0.28324415 |
| SH3PXD2A | -2.1081526 | 0.00804668 | 0.28324415 |
| TPM2 | -3.1066162 | 0.00806861 | 0.28324415 |
| HAPLN3 | -2.9525821 | 0.00807205 | 0.28324415 |
| RDH10-AS1 | 3.58104646 | 0.00812532 | 0.28389519 |
| GAS6 | -2.1746006 | 0.00814054 | 0.28389519 |
| PARM1 | -2.98433 | 0.00831625 | 0.28913578 |
| ALCAM | 2.16806768 | 0.0084077 | 0.2914241 |
| MYL9 | -2.4313219 | 0.00848884 | 0.29334222 |
| ENG | -2.265635 | 0.00864398 | 0.29779834 |
| HEYL | 2.62173496 | 0.00872741 | 0.29889337 |
| SCUBE3 | -3.4314166 | 0.00874401 | 0.29889337 |
| HHIP | 3.47444436 | 0.00875464 | 0.29889337 |
| EFEMP2 | -2.2590874 | 0.00879623 | 0.29941417 |
| RAPGEF5 | -3.0925829 | 0.00894085 | 0.30342847 |
| SOX11 | 3.76732919 | 0.00916661 | 0.30935523 |
| ZNF702P | 2.90599506 | 0.00918583 | 0.30935523 |
| TNFRSF21 | 2.57765329 | 0.00919712 | 0.30935523 |
| PIANP | -2.7139502 | 0.00924672 | 0.31010591 |
| FANK1 | -2.9918277 | 0.00929635 | 0.31085363 |
| SEC14L2 | -2.0921018 | 0.00933216 | 0.31113591 |
| GREM2 | -5.1250482 | 0.00939542 | 0.312329 |
| PODXL | 2.15602942 | 0.00950564 | 0.3150718 |
| FZD8 | -2.2238347 | 0.00991395 | 0.32765036 |
| LOC1019283( | 4.16502189 | 0.01023159 | 0.33619356 |
| SVIP | 1.9688869 | 0.01051897 | 0.34071266 |
| AQP3 | 3.05325008 | 0.010797 | 0.34480659 |
| CLDN19 | 4.84959988 | 0.01089748 | 0.34510703 |
| VAT1L | 3.6492856 | 0.01155907 | 0.3564127 |
| DEPDC1 | 3.08451383 | 0.01160504 | 0.3564127 |
| TMEM215 | 4.04465667 | 0.01217665 | 0.36720525 |
| NEDD1 | 1.80186651 | 0.01235725 | 0.37166548 |
| CD22 | 3.29117039 | 0.0127876 | 0.37940406 |
| CENPI | 2.73492427 | 0.01300117 | 0.38292828 |
| PLEKHG5 | 2.13617229 | 0.0135704 | 0.39493497 |
| KIF19 | 4.99583055 | 0.01378031 | 0.39864729 |
| MMS22L | 2.32575955 | 0.01407288 | 0.40504956 |

| | | | |
|---|---|---|---|
| TMPRSS5 | 3.10449084 | 0.01424243 | 0.40786454 |
| LRRC8B | 2.31109002 | 0.01502469 | 0.42281121 |
| CAMK2N1 | 1.83720677 | 0.01519977 | 0.42668192 |
| E2F7 | 3.00371544 | 0.0153046 | 0.42856653 |
| ARHGAP29 | 2.21351865 | 0.01560729 | 0.43383691 |
| SHROOM2 | 2.17372466 | 0.0162677 | 0.44733106 |
| LOC730101 | 1.93437872 | 0.01628948 | 0.44733106 |
| CDCA7 | 2.72633696 | 0.01641476 | 0.44968529 |
| POU3F2 | 3.25283137 | 0.01658103 | 0.4509803 |
| PM20D2 | 1.96034368 | 0.01685549 | 0.45409976 |
| KCNQ1OT1 | 2.27747111 | 0.01719741 | 0.45788613 |
| LIPC | 3.61730027 | 0.01755427 | 0.46304988 |
| CHD1 | 1.6276701 | 0.01771573 | 0.4640786 |
| CACNA2D2 | 3.38928866 | 0.01781671 | 0.46458296 |
| PLAU | 1.90297788 | 0.01799676 | 0.46713501 |
| TSPAN11 | 3.34877475 | 0.01824513 | 0.47142923 |
| LAMA1 | 2.34701861 | 0.0185265 | 0.4745136 |
| CLSPN | 2.95728023 | 0.018557 | 0.4745136 |
| BUB1 | 2.64660869 | 0.01857926 | 0.4745136 |
| FZD6 | 1.91086787 | 0.01863783 | 0.4745136 |
| FGFBP2 | 3.81013533 | 0.01932622 | 0.48610575 |
| BARD1 | 2.1943747 | 0.01943578 | 0.48778242 |
| IL17B | 3.64302592 | 0.01971116 | 0.48849732 |
| CSPG4 | 1.87226752 | 0.02141892 | 0.51373766 |
| GRIA4 | 3.60751256 | 0.02187239 | 0.51900067 |
| DUSP4 | 1.68690915 | 0.02196522 | 0.51900067 |
| NCMAP | 5.11122568 | 0.02204982 | 0.51900067 |
| HOXC8 | 1.94923375 | 0.02209485 | 0.51900067 |
| PIEZO2 | 2.23494461 | 0.02220284 | 0.52030099 |
| ITIH3 | 2.78117855 | 0.02224173 | 0.52030099 |
| CDKN1C | 2.93899898 | 0.02239767 | 0.52287293 |
| SHISA7 | 3.55854333 | 0.02247476 | 0.52359746 |
| ATP8B1 | 2.66767684 | 0.02259751 | 0.5248906 |
| CARNS1 | 2.68240678 | 0.02263599 | 0.5248906 |
| RAPGEF6 | 1.76291756 | 0.02286589 | 0.52653641 |
| RIF1 | 1.70319517 | 0.0228709 | 0.52653641 |
| ECT2 | 2.24607329 | 0.02330995 | 0.53520228 |
| RFC3 | 1.94264207 | 0.02334949 | 0.53520228 |
| SHCBP1 | 2.49748687 | 0.0236118 | 0.5381093 |
| S1PR1 | 2.45816827 | 0.02361831 | 0.5381093 |
| C18orf54 | 2.36730204 | 0.02376292 | 0.54026797 |
| HIST1H1C | 2.81028494 | 0.0238081 | 0.54026797 |
| SPINT1 | 2.70001318 | 0.02436432 | 0.54851071 |
| HEATR1 | 1.79968599 | 0.02442021 | 0.54868256 |
| KLHL11 | 1.92743927 | 0.02482327 | 0.55554288 |

| | | | |
|---|---|---|---|
| ARHGAP11A | 2.31012081 | 0.02515869 | 0.56194334 |
| BRCA1 | 1.94383196 | 0.02571307 | 0.57096082 |
| NCEH1 | 2.29195165 | 0.02623986 | 0.5815224 |
| CDK6 | 1.60371676 | 0.02640086 | 0.5839522 |
| SOX2 | 1.88694039 | 0.02696558 | 0.59399663 |
| LIN7C | 1.8054929 | 0.02723038 | 0.59649737 |
| COL13A1 | 2.52416322 | 0.02808892 | 0.60943299 |
| MPP6 | 1.65549295 | 0.02850492 | 0.61610725 |
| RAD18 | 1.89494121 | 0.02860736 | 0.61627705 |
| RBP1 | 3.06496479 | 0.02870585 | 0.61647897 |
| NDC1 | 1.64492762 | 0.02911337 | 0.61867269 |
| DOCK3 | 2.21026879 | 0.0292815 | 0.61992805 |
| CUBN | 2.58446556 | 0.02963609 | 0.62248905 |
| GRIK5 | 1.94448157 | 0.03022537 | 0.62508455 |
| WDHD1 | 2.0779125 | 0.03023982 | 0.62508455 |
| LCOR | 1.83769728 | 0.03134328 | 0.63975174 |
| FANCD2 | 2.10613239 | 0.03149384 | 0.64052493 |
| SACS | 1.94521067 | 0.03223614 | 0.65042114 |
| PVRL3 | 1.89205699 | 0.03229442 | 0.65042114 |
| RGS12 | 1.98018247 | 0.03271446 | 0.65105314 |
| KIAA1524 | 2.21262354 | 0.03271522 | 0.65105314 |
| CDK17 | 1.51054549 | 0.03275595 | 0.65105314 |
| AURKB | 2.05942651 | 0.03302874 | 0.6541878 |
| LRRC8C | 1.78155204 | 0.03311951 | 0.65484466 |
| XPO4 | 1.63396198 | 0.03349266 | 0.65992737 |
| MYEF2 | 1.55329433 | 0.0338787 | 0.66327489 |
| NUP155 | 1.7647764 | 0.03405504 | 0.66400008 |
| CASC5 | 2.73690681 | 0.03414393 | 0.66400008 |
| PTGDS | 2.66885172 | 0.03420663 | 0.66400008 |
| UTP20 | 1.61740785 | 0.03463507 | 0.66967026 |
| NEDD4L | 1.53043036 | 0.03496055 | 0.67473847 |
| TBC1D4 | 1.61009305 | 0.03506693 | 0.67473847 |
| KIF20B | 1.80727472 | 0.03522002 | 0.67473847 |
| TNFRSF25 | 2.37740906 | 0.03575229 | 0.68085047 |
| GFAP | 2.69980372 | 0.03602067 | 0.68465309 |
| CEP85 | 1.50225427 | 0.03682769 | 0.69435157 |
| MLIP | 3.939712 | 0.03701207 | 0.69591459 |
| TGFB2 | 1.77046952 | 0.0370661 | 0.69591459 |
| FXYD6 | 3.04422552 | 0.03731812 | 0.69591459 |
| PTPRK | 1.60346306 | 0.03731837 | 0.69591459 |
| NFE2L3 | 1.62096744 | 0.03857984 | 0.70973171 |
| RBL1 | 1.75536023 | 0.03873197 | 0.71025508 |
| CTPS2 | 1.57254719 | 0.03875672 | 0.71025508 |
| ABCA10 | 2.71194226 | 0.03928458 | 0.7146023 |
| HAUS6 | 1.65354261 | 0.03946484 | 0.71673437 |

| | | | |
|---|---|---|---|
| SLC38A1 | 2.4606614 | 0.03973965 | 0.72057436 |
| PHIP | 1.44501217 | 0.0398552 | 0.72147628 |
| ADAMTS9 | 2.32387832 | 0.04007556 | 0.72147628 |
| KDM6A | 1.49908083 | 0.04010602 | 0.72147628 |
| OXTR | 1.89894712 | 0.04010669 | 0.72147628 |
| EMB | 2.05221572 | 0.0404238 | 0.72374525 |
| PLK4 | 2.13520559 | 0.04155698 | 0.73937612 |
| CNKSR2 | 1.97015833 | 0.04228619 | 0.74467934 |
| ANLN | 2.22116844 | 0.04241942 | 0.74538857 |
| DGKH | 1.79480316 | 0.04267612 | 0.74874204 |
| TFAM | 1.54994088 | 0.04305293 | 0.75128078 |
| NR3C2 | 2.08733932 | 0.04335086 | 0.75248972 |
| ZNF100 | 1.9521201 | 0.04336491 | 0.75248972 |
| STXBP5 | 1.48002785 | 0.04341923 | 0.75248972 |
| AGO2 | 1.96834505 | 0.04372185 | 0.75289898 |
| MCTP1 | 2.07237264 | 0.04393436 | 0.75289898 |
| RASA2 | 1.60430818 | 0.04405012 | 0.75289898 |
| MED12L | 2.32818831 | 0.04409768 | 0.75289898 |
| PGBD5 | 2.57611485 | 0.04415932 | 0.75289898 |
| CCNJL | 1.89010174 | 0.04486899 | 0.76136646 |
| NAA15 | 1.48942088 | 0.04515461 | 0.76393275 |
| ERC2 | 3.08367965 | 0.04602316 | 0.77126235 |
| SMC4 | 1.58902107 | 0.04606273 | 0.77126235 |
| MCM8 | 1.85023792 | 0.04626617 | 0.77352961 |
| PP7080 | 1.73752554 | 0.04642261 | 0.77500538 |
| EIF1AX | 1.3764922 | 0.04696494 | 0.77745196 |
| TRIM59 | 2.28937883 | 0.04704784 | 0.77745196 |
| CEP55 | 2.14088375 | 0.04743397 | 0.78120204 |
| LRP6 | 1.31682039 | 0.04754731 | 0.78120204 |
| FGF5 | 3.1468526 | 0.04754964 | 0.78120204 |
| SERPINE1 | 2.35622078 | 0.04778182 | 0.7819543 |
| NT5DC3 | 1.50449875 | 0.04780176 | 0.7819543 |
| NAA25 | 1.44885476 | 0.04793946 | 0.78308008 |
| KIF11 | 2.02460286 | 0.0483277 | 0.78622904 |
| EPG5 | 1.57232797 | 0.0483397 | 0.78622904 |
| CREB5 | 1.92922325 | 0.04918928 | 0.79155068 |
| LONRF2 | 2.53770539 | 0.04954908 | 0.7952877 |
| PP12613 | 2.99353677 | 0.04974349 | 0.79652639 |
| P2RY1 | 2.77330355 | 0.0499947 | 0.79836558 |

**Table S3.** **Differentially expressed genes in tumor-associated macrophage after anti-IL-6 treatment**

| | Gene | baseMean | log2FoldCh | lfcSE | stat | pvalue | padj |
|---|---|---|---|---|---|---|---|
| 1 | FOS | 439.0406 | -0.85473 | 0.193322 | -4.42126 | 9.81E-06 | 0.010372 |
| 2 | DUSP1 | 281.1922 | -0.80209 | 0.19564 | -4.09985 | 4.13E-05 | 0.012202 |
| 3 | ATF3 | 120.635 | -0.99247 | 0.2436 | -4.07418 | 4.62E-05 | 0.012202 |
| 4 | CXCL2 | 84.46732 | -1.04046 | 0.252938 | -4.11351 | 3.9E-05 | 0.012202 |
| 5 | KLF6 | 187.9575 | -0.69842 | 0.179971 | -3.8807 | 0.000104 | 0.017694 |
| 6 | FOSB | 96.66998 | -0.98712 | 0.256264 | -3.85195 | 0.000117 | 0.017694 |
| 7 | CTSK | 139.3682 | 0.95153 | 0.245323 | 3.878687 | 0.000105 | 0.017694 |
| 8 | JUN | 92.87043 | -0.83645 | 0.221201 | -3.78142 | 0.000156 | 0.020603 |
| 9 | MT-RNR2 | 3658.146 | -0.51575 | 0.141709 | -3.63949 | 0.000273 | 0.032083 |
| 10 | EGR1 | 226.2903 | -0.63282 | 0.180101 | -3.51367 | 0.000442 | 0.046716 |
| 11 | PPP1R15A | 48.73857 | -0.93597 | 0.282049 | -3.31846 | 0.000905 | 0.086978 |
| 12 | ZFP36 | 151.0349 | -0.8313 | 0.252524 | -3.29198 | 0.000995 | 0.087628 |
| 13 | HSPA5 | 516.4567 | -0.4752 | 0.151599 | -3.13457 | 0.001721 | 0.139937 |
| 14 | EIF5 | 65.16305 | -0.74138 | 0.252379 | -2.93756 | 0.003308 | 0.249762 |
| 15 | PSAP | 1107.952 | -0.52455 | 0.200089 | -2.62156 | 0.008753 | 0.56627 |
| 16 | CCNL1 | 88.01537 | -0.59229 | 0.223714 | -2.64751 | 0.008109 | 0.56627 |
| 17 | PSMD8 | 57.7257 | -0.73821 | 0.283056 | -2.60799 | 0.009107 | 0.56627 |
| 18 | AXL | 50.49142 | 0.780538 | 0.302473 | 2.580518 | 0.009865 | 0.579307 |
| 19 | SOCS3 | 76.36558 | -0.6589 | 0.257268 | -2.56116 | 0.010432 | 0.580361 |
| 20 | M6PR | 97.08885 | -0.50304 | 0.215969 | -2.32921 | 0.019848 | 0.667539 |
| 21 | SDC4 | 75.57372 | -0.54663 | 0.243288 | -2.24684 | 0.024651 | 0.667539 |
| 22 | H13 | 82.36299 | 0.571095 | 0.254165 | 2.246948 | 0.024643 | 0.667539 |
| 23 | TNS3 | 176.2372 | 0.422303 | 0.18998 | 2.222882 | 0.026224 | 0.667539 |
| 24 | PER1 | 63.13003 | -0.5898 | 0.265204 | -2.22396 | 0.026151 | 0.667539 |
| 25 | CTSB | 845.5958 | -0.41087 | 0.172185 | -2.38621 | 0.017023 | 0.667539 |
| 26 | PLEC | 1036.672 | 0.468022 | 0.2001 | 2.338937 | 0.019339 | 0.667539 |
| 27 | PDIA4 | 61.7903 | -0.56802 | 0.256042 | -2.21844 | 0.026525 | 0.667539 |
| 28 | FN1 | 172.7414 | 1.545741 | 0.672375 | 2.298926 | 0.021509 | 0.667539 |
| 29 | RGS1 | 129.3138 | -0.48066 | 0.203688 | -2.35979 | 0.018285 | 0.667539 |
| 30 | PFKFB3 | 59.97299 | -0.57719 | 0.254926 | -2.26416 | 0.023565 | 0.667539 |
| 31 | GPNMB | 151.5311 | -1.07865 | 0.4541 | -2.37535 | 0.017532 | 0.667539 |
| 32 | FLNA | 592.0818 | 0.380643 | 0.167467 | 2.27294 | 0.02303 | 0.667539 |
| 33 | TXNIP | 144.4615 | 0.458005 | 0.200055 | 2.289393 | 0.022057 | 0.667539 |
| 34 | IER2 | 61.80011 | -0.58399 | 0.261139 | -2.23632 | 0.025331 | 0.667539 |
| 35 | DPEP2 | 74.08311 | 0.660161 | 0.268298 | 2.460548 | 0.013872 | 0.667539 |
| 36 | SERPINB6A | 46.52316 | 0.669583 | 0.29052 | 2.304778 | 0.021179 | 0.667539 |
| 37 | MT-ND2 | 405.5573 | -0.36503 | 0.152061 | -2.40052 | 0.016372 | 0.667539 |
| 38 | MT-CO1 | 2980.569 | -0.3527 | 0.143073 | -2.4652 | 0.013694 | 0.667539 |
| 39 | MT-ND5 | 936.5215 | -0.34475 | 0.146331 | -2.35596 | 0.018475 | 0.667539 |
| 40 | MT-ND6 | 166.7469 | -0.45694 | 0.202629 | -2.25507 | 0.024129 | 0.667539 |
| 41 | MT-CYTB | 700.027 | -0.35879 | 0.154906 | -2.31615 | 0.02055 | 0.667539 |
| 42 | GM13050 | 47.72761 | 0.642661 | 0.283059 | 2.270409 | 0.023183 | 0.667539 |
| 43 | CD9 | 53.39522 | 0.602215 | 0.277923 | 2.166841 | 0.030247 | 0.743514 |
| 44 | TAPBP | 118.0162 | -0.50097 | 0.240187 | -2.08575 | 0.037001 | 0.888869 |
| 45 | RIN2 | 59.46545 | 0.577632 | 0.280559 | 2.058865 | 0.039507 | 0.907807 |

| | | | | | | | |
|---|---|---|---|---|---|---|---|
| 46 | CALR | 322.9139 | -0.33921 | 0.164588 | -2.06096 | 0.039307 | 0.907807 |
| 47 | BTG2 | 90.45871 | -0.499 | 0.243853 | -2.0463 | 0.040727 | 0.915919 |
| 48 | OGT | 100.3106 | 0.438686 | 0.216499 | 2.026272 | 0.042737 | 0.941103 |
| 49 | ITM2B | 114.2228 | -0.41813 | 0.209893 | -1.99213 | 0.046357 | 0.98462 |
| 50 | PLIN2 | 62.33048 | -0.51003 | 0.256277 | -1.99013 | 0.046576 | 0.98462 |

**Table S4. Primer sequences used in qRT-PCR assay.**

| Genes | $T_m$ ($^{o}$C) | Forward Primer | Reverse Primer |
|---|---|---|---|
| *muTlr4* | 58 | TTC CTG CTG TTT CTC TTA CAC CT | GAG GCC AAT TTT GTC TCC ACA |
| *muCcl2* | 54.1 | GTC TGT GCT GAC CCC AAG AAG | TGG TTC CGA TCC AGG TTT TTA |
| *muIl6* | 58 | TAG TCC TTC CTA CCC CAA TTT CC | TTG GTC CTT AGC CAC TCC TTC |
| *muIl1β* | 58 | TGC CAC CTT TTG ACA GTG AT | TGT CCT CAT CCT GGA AGG TC |
| *muIfnγ* | 56.8 | CCA GCT GTT GCC GGA ATC | CGA ATC AGC AGC GAC TCC TT |
| *muTnfα* | 59.3 | CGG ACA GCT CGC TCT GCT A | TCC AGA GCC TGG CAC ACA |
| *muIfnα* | 62.7 | ATG GCT AGG CCC TTT GCT TTC | CAG TTC CTT CAT CCC GAC CAG |
| *muIfnβ* | 60.9 | CAG CTC CAA GAA AGG ACG AAC | GGC AGT GTA ACT CTT CTG CAT |
| *muVegfa* | 62.9 | CGG GAT TGC AGG GAA ACT | GGA TAC TAC GGA GCG AGA AG AG |
| *muGapdh* | 57.6 | AGG TCG GTG TGAACG GAT TTG | TGT AGA CCA TGT AGT TGA GGT CA |
| *muActin* | 57.6 | GGC TGT ATT CCC CTC CAT CG | CCA GTT GGT AAC AAT GCC ATG T |
| *muIl33* | 55.1 | TCC AAC TCC AAG ATT TCC CCG | CAT GCA GTA GAC ATG GCA GAA |
| *muCox2* | 54.8 | TGA GCA ACT ATT CCA AAC CAG C | GCA CGT AGT CTT CGA TCA CTA TC |
| *muNlrp3* | 57.9 | ATT ACC CGC CCG AGA AAG G | TCG CAG CAA AGA TCC ACA CAG |
| *muHmgb1* | 55.7 | GCT GAC AAG GCT CGT TAT GAA | CCT TTG ATT TTG GGG CGG TA |
| *huHMGB1* | 60.7 | TAT GGC AAA AGC GGA CAA GG | CTT CGC AAC ATC ACC AAT GGA |
| *huIL6* | 60.9 | CCT GAA CCT TCC AAA GAT GGC | TTC ACC AGG CAA GTC TCC TCA |
| *huIFNγ* | 57.9 | TCGGTAACTGACTTGAATGTCCA | TCGCTTCCCTGTTTTAGCTGC |
